# Semantic ambiguity dissociates the roles of word meaning and co-occurrence statistics in sentence processing

**DOI:** 10.64898/2026.05.15.724229

**Authors:** Andrey Zyryanov, Victoria Pierz, Yulia Oganian

## Abstract

Next-word prediction is a key component of human language processing, which manifests itself in higher processing costs (longer reading times and enhanced neural responses) to unpredictable words. These costs are captured by large language model (LLM) surprisal, a metric inferred from statistical patterns of word co-occurrence. This makes word co-occurrence statistics a plausible candidate source for human next-word prediction. In contrast, psycholinguistic models posit that human next-word prediction relies on representations of word meaning. To contrast these views, we placed words that carry multiple meanings in contexts that rendered meaning ambiguous (e.g., ‘My colleagues portrayed the *head’*: body part or leader). With this, we kept word co-occurrence statistics available for prediction but introduced uncertainty into word meaning, effectively decorrelating these putative drivers of next-word prediction. Across three self-paced reading and magnetoencephalography (MEG) experiments, human processing costs for unpredictable unambiguous words were captured by LLM word surprisal, while ambiguity disrupted this relation. This shows that human next-word prediction is mechanistically linked to ambiguity resolution, diverging from LLM-like statistics-driven prediction specifically under ambiguity. We further found that a shared cortical network supports prediction and ambiguity resolution in parallel. Overall, our findings corroborate a distinctive feature of predictive language processing in humans, namely its reliance on word meaning beyond statistical patterns of word co-occurrence.

**Significance Statement:** Next-word prediction is a key principle of human language comprehension, and the performance goal of large language models (LLMs). By design, LLMs exploit statistical patterns of word co-occurrence to predict upcoming words. In contrast, the basis for human next-word prediction remains debated. To understand whether word co-occurrence statistics likewise underlie prediction in humans, we designed sentences with unpredictable ambiguous words, such as ‘My colleague portrayed the head [leader vs. body part]’. LLM predictions captured how humans processed these words when context disambiguated their meaning but not when word meaning remained ambiguous. This reveals a distinctive feature of predictive language processing in humans: It is contingent on word meaning beyond statistical regularities of word usage.

## Introduction

To comprehend language, humans infer sentence meaning in a predictive fashion (Altmann and Kamide, 1999; Federmeier, 2007; Lau et al., 2013; Kuperberg and Jaeger, 2016; Pickering and Gambi, 2018; Staub, 2025). In a key step of this process, the brain evaluates the mismatch between predicted and perceived word representations, forming the lexico-semantic prediction error (Fig. 1a, b; Bornkessel-Schlesewsky & Schlesewsky, 2019; Fitz & Chang, 2019; Nour Eddine et al., 2024; Rabovsky et al., 2018). Greater prediction errors for unexpected words are reflected in longer reading times (Smith and Levy, 2013; Wilcox et al., 2023; Shain et al., 2024; Staub, 2025) and enhanced N400 neural responses (Kutas and Hillyard, 1980; Helenius et al., 1998; Frank et al., 2015; Nieuwland et al., 2020; Heilbron et al., 2022; Zou et al., 2026). However, the nature of word representations necessary for prediction error computation remains unclear.

**Figure 1.**
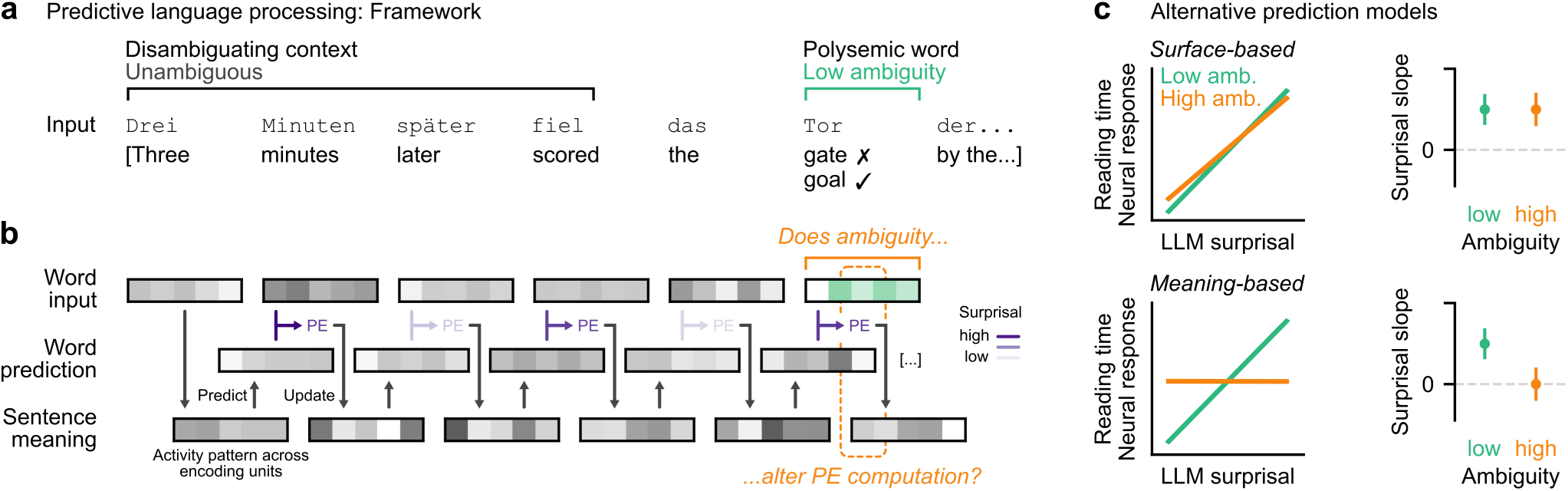
Predictive language processing: Framework and alternative hypotheses. (a) An example sentence with a polysemic word in a disambiguating context. (b) Three key stages of predictive language processing. *Top row:* Each word input is represented in a distributed pattern of neural activity (representation vector shown schematically in grey shades). *Middle row:* Predictions of upcoming words in the same representational space as for word inputs. *Bottom row:* Evolving representation of the entire sentence. *Predict:* The current representation of sentence meaning generates a prediction of the upcoming word. *Prediction error (PE):* When the word is perceived, its representation is compared against the prediction, generating the lexico-semantic prediction error. *Update:* This prediction error governs the update of the sentence meaning representation. We asked whether the lexico-semantic prediction error is based on co-occurrence statistics of word forms (surface statistics) or word meaning. To answer, we leveraged polysemy to selectively introduce uncertainty into the meaning representation of the word perceived. If PE computation relies on word meaning rather than surface statistics, we expect to find that behavioral and neural signatures of PE computation will be altered by this uncertainty. (c) The effects of surprisal and ambiguity on reading times and neural responses expected under surface-versus meaning-based prediction models. Word surprisal from a large language model (LLM) served as a proxy of PE inferred from statistical patterns of word co-occurrences. Under the surface-based prediction model, we expected LLM surprisal to predict longer reading times and larger neural responses regardless of ambiguity. Under the meaning-based prediction model, we expected ambiguity to alter prediction error computation, disrupting the relation between LLM surprisal and either dependent variable. Of note, we made no specific predictions regarding the main effect of ambiguity: it differs between polysemic words with related versus unrelated meanings (Eddington and Tokowicz, 2015), whereas the meanings of the polysemic words used here were only partially related.

The rise of large language models (LLMs) has inspired the hypothesis that human language processing relies on statistical patterns of word co-occurrence (Casto et al., 2025). LLMs reach human-like performance in text production relying exclusively on such statistical patterns (Devlin et al., 2019; Radford et al., 2018). Crucially, LLM estimates of word surprisal capture prediction error signatures both in reading times (Smith and Levy, 2013; Wilcox et al., 2023; Staub, 2025) and in N400 responses (Goldstein et al., 2022; Heilbron et al., 2022; Zou et al., 2026). Moreover, the structure of LLM representations predicts neural activity patterns during language comprehension (Schrimpf et al., 2021; Caucheteux and King, 2022; Goldstein et al., 2022). This alignment motivates the hypothesis that lexico-semantic prediction error is surface-based, increasing for words that occur in statistically atypical contexts regardless of how well they fit such contexts semantically.

The computational goal of human language processing, however, is to infer sentence meaning. Hence, psycholinguistic models posit that lexico-semantic prediction error increases whenever a word carries unexpected meaning (Rabovsky et al., 2018; Nour Eddine et al., 2022, 2024). Indeed, neural signatures of prediction error are sensitive to uncertainty in sentence meaning, for example due to limited preceding context (Zou et al., 2026), and to a word’s contribution to overall sentence meaning (Thornhill and Van Petten, 2012; Stone et al., 2023). However, words semantically unexpected in a particular context would also be statistically atypical in that context (Günther et al., 2019; Fernandino et al., 2022), conflating the surface-and meaning-based models. Distinguishing between them is essential to pinpoint the processing stage where surface representations of language input and its meaning are integrated (Casto et al., 2025).

We distinguished these models using words with multiple meanings (e.g., ‘bat’: animal or device). Such polysemic words are pervasive (Rodd et al., 2002; Zipf, 1949) and require the brain to resolve co-activation of multiple word meanings (Chen & Boland, 2008; Duffy et al., 1988, 2001; Rodd, 2020; Swaab et al., 2003). We manipulated sentential contexts before these words to vary uncertainty in their meaning, or ambiguity. We then tested how well behavioral and neural signatures of prediction error were modelled by LLM word surprisal under low and high ambiguity (Fig. 1c). We hypothesized that, if next-word prediction is surface-based, uncertainty in word meaning should not influence prediction error computation. In contrast, if it is meaning-based, ambiguity should introduce uncertainty into prediction error computation and thereby alter its behavioral and neural signatures.

This setup also allowed us to directly dissociate prediction error computation and ambiguity resolution. When ambiguity resolution is hampered by insufficient context, uncertainty in word meaning enhances neural responses around 400 ms after word onset (Beretta et al., 2005; Lee and Federmeier, 2006; Pylkkänen et al., 2006; Ihara et al., 2007; Maciejewski and Klepousniotou, 2020), in temporal overlap with the N400 signatures of prediction error. However, it is unknown how the two processes interact. By decorrelating measures of ambiguity and prediction error in our stimuli, we tested whether they rely on the same brain networks, are mechanistically linked, or proceed independently.

## Materials and Methods

### Participants

#### Online self-paced reading experiments

In Experiment 1, 150 participants (88 males, 54 females, 5 non-binary, 3 preferred not to respond; mean age 32.2 years, range 18 to 50 years; 128 right-handed) read sentences from one of three non-overlapping experimental lists, yielding 47 to 52 participants per sentence. In Experiment 2, 55 participants (41 male, 13 female, 1 non-binary; mean age 32.9 years, range 18 to 49 years; 46 right-handed) read sentences from a single experimental list. Of these participants, 19 (35%) also took part in Experiment 1. The two experiments were conducted 7 months apart, ruling out potential learning effects. Participants were recruited via Prolific and were reimbursed at a rate of €12 per hour.

All participants were native speakers of German, and none reported dyslexia or language disorders. Most participants (76% in Experiment 1 and 62% in Experiment 2) reported normal vision; the remaining participants did not differ from those who reported normal vision in task performance (rating correlation: Pearson *r* = 0.98, *p* < 0.001).

#### Magnetoencephalography (MEG) experiment

27 participants (12 males, 13 females, 2 non-binary; mean age 25.1 years, range 20 to 40 years; all right-handed) completed the MEG experiment. All were native speakers of German raised in a monolingual setting. All reported normal or corrected-to-normal vision and normal hearing; none reported dyslexia or language disorders. Participants were recruited via a university mailing list and were reimbursed at a rate of €15 per hour.

#### Ethics approval

The study was approved by the institutional review board of the Medical Faculty of the University of Tübingen (No. 728/2023BO2). All participants provided informed consent before starting the study. All experiments were conducted in accordance with the Declaration of Helsinki.

### Online Experiment 1

#### Stimuli

We created 300 grammatical German sentences with polysemic target words: half contained the word ‘Blatt’ (meaning ‘paper’ or ‘leaf’), the others contained ‘Tor’ (meaning ‘gate’ or ‘goal’). The target words were comparable in word form frequency (Blatt: 12.17 instances per million tokens, ipm, Tor: 48.9 ipm; retrieved from SUBTLEX-DE, Brysbaert et al., 2011) and were matched in number and case across sentences (Blatt/Tor counts: 74/76 singular, 76/74 plural; case: 52/52 nominative, 48/48 dative, 50/50 accusative).

Target words were selected based on a previous norming study (Moritz et al., 2001), such that the two meanings of each target word were roughly balanced in frequency. Moritz and colleagues (2001) asked 100 native speakers of German to provide one verbal association to each polysemic word. These associations were assigned to one of the possible meanings for each word. Of all responses, 93–95% referred to one of the two meanings employed in the present study. Of these responses, 57% referred to the dominant meaning (‘leaf’ or ‘gate’), while the remaining 43% referred to the other meaning (‘paper’ or ‘goal’). Given roughly balanced meaning frequencies, sentential context was the primary source of disambiguating information.

A native speaker of German trained in linguistics created pre-target contexts (mean length 6 words, range 5 to 9 words) to constrain target interpretation towards one of its meanings or keep it ambiguous. The post-target context was a grammatical and semantically plausible completion of the sentence. The post-target context served to avoid an overlap between prediction error signatures and auditory offset responses in subsequent MEG recordings. However, since our research question focused on target processing, we manipulated disambiguating information only before the target word. In five of 300 sentences, we noticed minor typing errors after collecting the data of Experiment 1. These sentences were excluded from Experiment 1 analyses but were corrected for further experiments.

#### Procedure

The sentences were split into three experimental lists matched in Blatt/Tor counts; each participant was presented with a single list. Participants’ task was to read each sentence and to indicate what the last word meant in this sentence. The sentences were truncated after the target word. Before each trial, participants were shown whether the upcoming sentence will finish with ‘Blatt’ or ‘Tor’. During the trial, participants had to read the sentence word-by-word in a self-paced manner by pressing the space key. The words were presented in the center of the screen, one at a time, with their adjacent punctuation. At the end of the trial, participants moved a slider towards one of the word meanings depicted with emojis at the extremes of the slider. The midpoint was labelled as ‘unklar’ (unclear). The initial slider position was always at the midpoint. Participants were instructed that the emojis did not represent specific meanings, but rather prototypical ones: for example, if a sentence referred to the Brandenburg Gate, they had to move the slider towards the emoji of the gate even though it did not depict the Brandenburg Gate exactly. The target words never repeated on consecutive trials to mitigate direct meaning priming. The experiment was hosted on Pavlovia (pavlovia.org) with presentation controlled by PsychoJS (version 2022.2.5).

### Online Experiment 2

#### Stimuli

Stimuli were 400 grammatically valid sentences. Half of the sentences contained a semantically anomalous word (for example, ‘Fein samtig überzogen sind die *Irrtümer* der Sukkulenten.’: ‘Covered with a fine, velvety coating are the *mistakes* of the succulents’). The non-anomalous sentences comprised a subset of 100 original sentences from Experiment 1 where target ambiguity and surprisal (see *Stimulus features* below) were orthogonal (henceforth, AS-orthogonal sentences; *N* = 50 per target word; see *Results* and Fig. 2c), and new filler sentences (*N* = 100) that did not contain either of the polysemic target words. The semantically anomalous sentences were the remaining sentences from Experiment 1 (*N* = 200) where one word was replaced either at the target position (*N* = 101) or at an arbitrary position after the target (*N* = 99). Thus, each sentence was equally likely to contain one of the polysemic target words or a semantic anomaly. Furthermore, the sentences that did not contain the polysemic target words contained one of four other words: ‘Ferien’ (holiday), ‘Wein’ (wine), ‘Gesetz’ (law), ‘Irrtum’ (error). Thus, participants could expect some words to repeat across trials but were blind to which words were essential to our research question. The sentences contained an average of 11.9 words (range 5–18 words).

**Figure 2.**
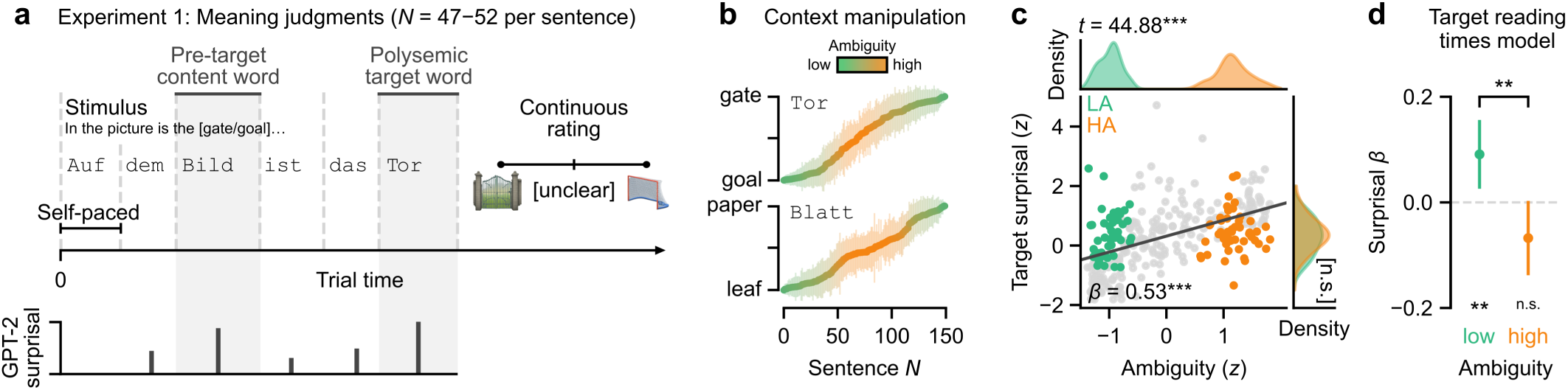
The relation between GPT-2 surprisal, ambiguity, and reading times. (a) Single trial setup in Experiment 1. Participants read sentences on the screen word by word in a self-paced manner up to and including the polysemic target word. Each sentence contained 6 to 9 words before the target word. After the target word, participants rated its perceived meaning on a continuous analog scale. (b) Ratings of target word (‘Blatt’ or ‘Tor’) meanings for each sentence (*N* = 150 per target word), averaged across participants and sorted in increasing order. Error bars denote standard deviation across participants. While some sentence contexts strongly favored one of two meanings (green, low ambiguity), others left the target word ambiguous (orange). We quantified target ambiguity as the negative absolute rating of target meaning for each sentence for all subsequent analyses. (c) The relation between target ambiguity and GPT-2 surprisal (dark grey line shows linear regression fit; adjusted *R^2^* = 0.26). Green and orange dots denote our selection of two sentence subsets that differed in target ambiguity but were matched on target surprisal (AS-orthogonal sentences; HA – high ambiguity; LA – low ambiguity). *Top:* Non-overlapping distributions of target ambiguity in the two subsets. *Right:* Overlapping distributions of target surprisal in the two subsets. Any differences in the effects of GPT-2 surprisal on reading times between the HA and LA subsets must be due to target ambiguity, as they are matched on target surprisal. (d) *β* coefficients for target surprisal based on a linear mixed-effects regression of log-transformed target reading times for AS-orthogonal sentences. Error bars represent 95% confidence intervals. ** *p* < 0.01, *** *p* < 0.001, n.s. – not significant.

#### Procedure

All 400 sentences were presented to each participant. Participants’ task was to read each sentence word-by-word and to rate how likely it was that the sentence came from an online blog, a book, or from others’ speech (one joint rating per sentence). In contrast to Experiment 1, they were not instructed to rate a particular word, but rather to look out for any words that rendered the sentence implausible and rate the sentence overall. The words were presented in the center of the screen, one at a time, with their adjacent punctuation. The extremes of the slider were labelled as ‘sehr unwahrscheinlich’ (very unlikely) and ‘sehr wahrscheinlich’ (very likely). The initial slider position was random. The polysemic words never repeated on consecutive trials. The experiment was hosted at Pavlovia (pavlovia.org) with presentation controlled by PsychoJS (version 2022.2.5).

### MEG experiment

#### Stimuli

Stimuli were the 300 sentences of Experiment 1, spoken in either a male or a female voice (*N* = 150 each). The audios were generated with a text-to-speech model (https://voicemaker.in/). We truncated the sentences by fading their volume to zero from, on average, 375 ms after target offset (range 300–450 ms). This allowed us to shorten stimulus duration and maximize the number of trials while avoiding auditory offset responses.

Word boundaries were annotated using a forced aligner (WebMAUS General; Kisler et al., 2017) and manually corrected for all words included in the MEG analyses. Average stimulus duration was 3559 ms (SD 572 ms, range 2437 to 5680 ms). Target onset was on average at 2238 ms (SD 549 ms, range 1255 to 4273 ms).

#### Procedure

Participants’ head shapes and the locations of three head position indicator (HPI) coils were digitized using a Polhemus 3SPACE Fastrak system (Polhemus, Vermont, USA). Participants were then seated in a dimly lit magnetically shielded room in a reclined position. After positioning their head in the MEG system, they were instructed that they would hear sentences that contain one of two words, ‘Blatt’ or ‘Tor’. Their task was to listen attentively to each sentence. Whenever a sentence was followed by two emojis that represented different target word meanings, participants were asked to choose the one that best fits the last sentence by pressing either the left or the right button. The task consisted of four 15-minute blocks with self-paced breaks in between.

The stimuli were presented binaurally via insert headphones (E-A-RTONE 3A) with loudness adjusted individually to each participant’s comfort. Consecutive sentences were separated by a silent interval (mean 500 ms, range 450–550 ms, sampled from a uniform distribution). During passive listening, a fixation cross was projected onto the screen. In catch trials (*N* = 32 in 19 participants; *N* = 80 in 8 participants), response options were projected onto the screen until a button was pressed. Task presentation was controlled by PsychoPy (version 2021.2.3).

Each sentence was repeated twice for a total of 600 trials. Their order was pseudo-randomized under two constraints: 1) repetitions of any given sentence were separated by at least 50 other sentences; 2) the target words never repeated in consecutive trials. The randomization was performed using Mix (van Casteren and Davis, 2006).

### Stimulus features

#### Ambiguity

Target ambiguity was quantified for each sentence based on the meaning ratings obtained in Experiment 1. Single-trial ratings on a scale between 0 and 1 were logit-transformed and averaged across participants for each sentence. Ambiguity was defined as the negative absolute value of this average rating and *z*-scored across sentences. Its maximum value corresponds to the midpoint of the slider, and decreases as slider positions approach any endpoint (that is, any particular meaning).

#### Plausibility

Plausibility was defined as the average sentence rating obtained in Experiment 2 and *z*-scored across sentences. Note that because the sentences in Experiment 2 continued after the target word, plausibility was affected by information both preceding and following the target word.

#### Surprisal

We passed each stimulus sentence through the GPT-2 model pre-trained on a German text corpus of ∼2 billion tokens and available at HuggingFace (https://huggingface.co/dbmdz/german-gpt2). The model consecutively processed each token in the sentence and predicted the next token given prior context. These predictions formed a probability distribution over all tokens in model vocabulary.

To compute word surprisal, we averaged the negative log-probability across all tokens contained in the model representation of each word. Note that the polysemic target words were complete tokens and never required averaging. In the online experiments, individual words were presented with their adjacent punctuation; therefore, we included punctuation tokens into averaging. One exception was the target word: since punctuation was presented visually in the reading experiments but not in the MEG experiment, we kept target surprisal values consistent between experiments by excluding punctuation tokens. Surprisal values with punctuation tokens were on average lower than those without (mean difference: –2.54 bits, SD 2.62 bits; paired *t*-test: *t*(470) = –21.05, *p* < 0.001), but the two were highly correlated (Pearson *r* = 0.94, *p* < 0.001).

We quantified target prediction uncertainty as Rényi entropies (Rényi, 1961) computed over the distribution of GPT-2 predictions at the target word position:

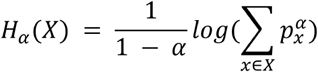

We computed this metric at the *α* orders of 0.1, 1, and 10. At *α* = 1, Rényi entropy is equivalent to Shannon entropy, a metric widely used in previous studies. The additional *α* values of 0.1 and 10 were motivated by a recent study that found Rényi entropy at these *α* values to optimally predict neural activity during speech processing (Guilleminot and Morillon, 2025).

Finally, we quantified how much GPT-2 predictions changed after versus before the target word was presented as the Kullback-Leibler divergence between their probability distributions. We included this metric to test whether less ambiguous words incur a stronger shift in model predictions.

#### Word frequency

Normalized word frequency was extracted from SUBTLEX-DE (Brysbaert et al., 2011; https://osf.io/py9ba). We added a constant of 1, log-transformed and *z*-scored these values.

#### Pre-target content word definition

Several behavioral and neural data analyses were conducted on the content word immediately preceding the target (henceforth, pre-target content word). These pre-target content words were any nouns, verbs, adjectives, or adverbs that immediately preceded the target (function words between the two were allowed) and were themselves preceded by two or more words. The latter was motivated by the finding that surprisal effects are attenuated at sentence onsets (Zou et al., 2026). These criteria yielded a total of 290 items in the pre-target content word analyses.

### Behavioral data preprocessing

We excluded trials with unrealistically short or long reading times (below 50 ms or above 2500 ms) and those exceeding 3 standard deviations above participant-specific means. This resulted in a total of 2.67% of trials excluded across Experiments 1 and 2. Reading times were log-transformed and *z*-scored for model fitting.

### MEG recording and preprocessing

#### Recording

Neural activity was recorded using a 275-sensor, whole-head CTF MEG system (VSM Medtech, Port Coquitlam, Canada) in a magnetically shielded room (Vakuumschmelze, Hanau, Germany). The sampling rate was 1171.875 Hz. Five continuously noisy sensors were disabled for all participants. Head position was continuously monitored via three head position indicator coils attached to the nasion, left and right pre-auricular points. Auditory stimuli were concurrently recorded by the MEG system for offline synchronization.

#### Preprocessing

All data were preprocessed and analyzed with MNE-Python (version 1.8.0; Gramfort et al., 2013) and custom Python scripts. The onset of each stimulus was identified by cross-correlating stimulus audios with those recorded by the MEG system. The data were notch-filtered at 50 and 100 Hz, band-pass-filtered between 0.5 and 20 Hz (both using the default MNE 1.8.0 settings), and downsampled to 300 Hz. In four participants, a single bad channel was interpolated. Heartbeat and ocular artefacts were removed using an independent component analysis.

Because target ambiguity and surprisal were reflected in the stimulus spectrogram (Fig. S2), we regressed the acoustic responses from neural activity using a spectro-temporal response function model (Holdgraf et al., 2017). The model was fitted to the full MEG time series of each participant, with each channel *z*-scored relative to its mean and standard deviation across the entire experiment. The predictors were stimulus spectrograms computed using a mel-scaled filter bank with 15 center frequencies spanning 50–5000 Hz. The spectrograms were *z*-scored within each frequency band, and silent inter-stimulus intervals were assigned the minimum value of that band. To account for neural response delays, we added time-lagged versions of the spectrogram with delays from 0 to 600 ms in ∼17 ms (5-sample) steps.

Model fitting was performed using ridge regression, with the regularization parameter selected via bootstrapping. Specifically, the data were randomly split into 80% training and 20% testing sets across 15 iterations. For each sensor, the optimal ridge parameter was chosen from 15 logarithmically spaced values from 10^−3^ to 10^8^ to maximize the average test-set correlation between predicted and observed neural activity. The model was then re-fitted to the full MEG time-series using the selected parameter, and its predictions were subtracted from the MEG signals to regress out variance explained by acoustic responses.

The residual neural responses were segmented from –250 to 750 ms around the onsets of the pre-target content word and of the target word. They were *z*-scored to the channel-specific mean and standard deviation computed over the silent interval from –250 to –50 ms before sentence onset across all trials.

#### Source reconstruction

The head shape and HPI coils digitized with Polhemus were aligned to the HPI coil locations recorded by the MEG system. Cortical, skull and scalp surfaces were reconstructed with FreeSurfer (Fischl, 2012) based on individual T1-weighted images acquired with a 3T Siemens Magnetom Prisma scanner (MP-RAGE sequence; repetition time 2300 ms; echo time 3 ms; voxel size 1 × 1 × 1 mm). These individual reconstructions (available for 25 participants) or the ‘fsaverage’ template (for the remaining 2 participants) were then aligned to the head shape digitization data in the MEG coordinate system.

To perform a minimum-norm-based reconstruction of source activity, we first defined an ico-4 source space on the white matter surface of each participant (2562 vertices per hemisphere). We computed the forward solution using a single-layer boundary element model with the default MNE 1.8.0 parameters. Noise covariance was estimated from empty room recordings conducted immediately before or after each recording session. The inverse solution was computed with source orientations fixed perpendicular to the cortical surface (Gwilliams et al., 2016). The inverse solution was applied to the average evoked responses or to single epochs (see below) with the signal-to-noise ratios set to three and two, respectively. This yielded the time-series of dSPM values for each source in each participant’s native anatomical space. Average responses or regression coefficients computed for each participant (see below) were warped into the common ‘fsaverage’ ico-4 source space for group-level visualization.

### Data analysis

#### Reading times

We fitted linear mixed-effects regression models to log-transformed and z-scored target reading times with the lmerTest package (version 3.1-3; Kuznetsova et al., 2017) in R (version 4.2.2).

In brief, all models included the following fixed effects of interest: GPT-2 surprisal of the word being analyzed, target ambiguity, and their interaction (Fig. 1c). We also included surprisal of two preceding words (referred to as surprisal at lags 1 and 2) to account for spill-over effects (Wilcox et al., 2023). Other covariates of no interest were: log-transformed word frequency, word length in characters, the interaction between word frequency and length, sentence position within trial list, word position within a sentence, the interaction between sentence and word position. For Experiment 2, the covariates additionally included sentence plausibility and its interaction with current word surprisal.

The initial random effect structure included by-item random intercepts, and uncorrelated by-participant random intercepts and random slopes for each fixed effect. We then eliminated random slopes with the lowest variance until the models converged without producing singular fit warnings. The full structure of each model is presented in Tables S1 – S3.

We first fitted models across all 300 sentences with ambiguity as a continuous variable. For follow-up analyses of the ambiguity-surprisal interaction, we restricted the models to AS-orthogonal sentences (*N* = 100; see *Results* and Fig. 2c) and entered ambiguity as a categorical treatment-coded variable with either high or low ambiguity as the baseline. This enabled us to estimate the effect of surprisal for each baseline level. All continuous predictors were *z*-scored.

#### Neural responses to ambiguity

We averaged sensor-space neural responses to low-and high-ambiguity target words in AS-orthogonal sentences per participant, and compared them across participants using a two-tailed spatiotemporal cluster permutation *t*-test. The cluster-forming threshold corresponded to a two-tailed *p*-value of 0.01. The maximum distance between samples to be considered temporally adjacent was set to 8 samples (27 ms), roughly half a cycle of neural activity at the upper frequency boundary of 20 Hz. The null distribution was computed over 5000 permutations. All subsequent (spatio-)temporal cluster permutation tests were conducted with the same parameters.

For an anatomical source-space visualization, we projected the average neural responses to the target word into the common ‘fsaverage’ space. For each timepoint and source, we computed the *t*-statistic comparing responses to the low-versus high-ambiguity target words across participants. We thresholded this statistic at ±2.05, which corresponds to an uncorrected two-tailed *p*-value of 0.05.

#### Sensor-space region of interest (ROI) definition

To maximize our sensitivity to the relation between surprisal and neural responses to the target word, we restricted the analyses to a sensor-space ROI. We defined the ROI based on neural responses to the pre-target content word under low target ambiguity. This made ROI definition orthogonal to our comparison of interest, namely, whether the relation between surprisal and neural responses differed between low-and high-ambiguity target words.

First, we regressed neural responses to the pre-target content word in the low-ambiguity subset of AS-orthogonal sentences against the GPT-2 surprisal of this word. The predictors also included preceding word’s surprisal as a covariate of no interest. The regressions were fitted separately for each timepoint, sensor, and participant. Next, we extracted the time series of *β* coefficients for surprisal in each sensor and subjected them to a principal component analysis (PCA). PCA was fitted in the time window from – 250 to 400 ms around pre-target content word onset to ensure that target-related activity did not affect PC weights. To test if the first PC reliably captured the N400 response, we compared its time series against zero with a two-tailed one-sample temporal cluster permutation *t*-test. Finally, we used its sensor weights as our spatial ROI.

#### Neural responses to surprisal as a function of ambiguity

We first examined how surprisal modulated N400 responses under low versus high ambiguity. We averaged single-trial neural responses to the target word over sensors using the spatial ROI weights, and over the *a priori* defined N400 window from 250 to 500 ms post-onset (Frank et al., 2015; Heilbron et al., 2022). We analyzed these responses with linear mixed-effects regression models with target ambiguity, GPT-2 surprisal, and their interaction as fixed effects (additional covariates of no interest were GPT-2 surprisal at lags 1 and 2; see Tables S4, S5 for the full model structure). The procedure for random slope selection was the same as in the reading times analyses.

To examine the effects of surprisal in a time-resolved manner, we regressed neural responses to the target word against target surprisal and preceding word’s surprisal (as a covariate of no interest). The regressions were fitted separately for each timepoint, sensor, participant, and ambiguity subset of AS-orthogonal sentences. We aggregated the *β* coefficients for target surprisal over sensors using the spatial ROI weights. We then compared *β* coefficients under low and high target ambiguity, as well as their difference, against zero using two-tailed one-sample temporal cluster permutation *t*-test.

For an anatomical source-space visualization of the ambiguity-surprisal interaction, we projected single-trial neural responses to the target word into each participant’s native source space. Next, we regressed neural responses in each source against target surprisal and preceding word surprisal. As above, this was done separately for the low-and high-ambiguity subsets of AS-orthogonal sentences. Next, we warped the time series of the *β* coefficients for target surprisal into the common ‘fsaverage’ template. Finally, we computed the *t*-statistic comparing the *β* coefficients between low-and high-ambiguity target words across participants, for each time point and source. We thresholded this statistic at ±2.05, which corresponds to an uncorrected two-tailed *p*-value of 0.05.

#### Topographical comparison of surprisal and ambiguity effects

The comparison between low-versus high-ambiguity target words yielded a participant-specific sensor-level topography of their difference at each time point in the target word window (matrix of shape # participants × # sensors × *#* time points). To quantify spatial alignment between ambiguity and surprisal effects over time, we computed cosine similarity between the spatial ROI of the surprisal effect and participant-specific topographies of the ambiguity effect at every time point, yielding a cosine similarity matrix of shape *#* participants × *#* time points. Finally, we compared cosine similarity against zero using two-tailed one-sample temporal cluster permutation *t*-test. A positive cosine similarity implies that the effect of ambiguity in individual participants is spatially aligned with the network generating surprisal responses.

#### The relation between ambiguity and prediction uncertainty

We examined how target ambiguity relates to the following metrics of uncertainty: Rényi entropies at the *α* orders of 0.1, 1 and 10, and Kullback-Leibler divergence (see above). First, we fitted a cross-validated linear regression of target ambiguity against all these metrics and target surprisal (full model). We quantified the unique variance each uncertainty metric explained in target ambiguity as the difference between test-set *R^2^* values of the full model versus the model where this metric was excluded. We then compared these *ΔR^2^* values against zero using upper-tailed one-sample *t*-tests. For this analysis, all variables were quantile-transformed, motivated by an observation that their relation is potentially non-linear, and *z*-scored.

For one metric that explained variance in target ambiguity (Rényi entropy at *α* = 0.1, or dispersion), we formally tested *post hoc* whether their relation was non-linear. We fitted two cross-validated linear regressions of target ambiguity against dispersion, one with and one without quantile-transforming both variables. Target surprisal was included into both models as a covariate of no interest. We compared the test-set *R^2^* between these models with a paired-samples two-tailed *t*-test.

These analyses were conducted on the 300 sentences of Experiment 1 and the MEG experiment. For cross-validation, the model was trained on 75% of the sentences and tested on the remaining ones. The procedure was repeated 50 times with random train-test splits to achieve stable *R^2^* estimates.

Finally, we addressed whether dispersion mediated the effects of target ambiguity on neural responses. We fitted a linear regression of neural responses to AS-orthogonal sentences against target ambiguity (categorical variable with low ambiguity as the baseline level), dispersion, their interactions with surprisal, and preceding word surprisal. The model was fitted separately for each sensor, time point, and participant. We aggregated the *β* coefficients for each predictor over sensors using the spatial ROI weights, and compared them against zero using two-tailed temporal cluster permutation *t*-test. This analysis was conducted first on the pre-target content word to test if its power was sufficient to detect potential effects of dispersion on neural responses. We then performed this analysis on the target word to test if dispersion mediated the main effect of ambiguity and the ambiguity-surprisal interaction.

#### Pre-target content word analyses

To address whether the interaction between ambiguity and surprisal was specific to target word processing, we fitted linear mixed-effects models to log-transformed reading times of the pre-target content word. Model structure repeated the target reading times analyses (Tables S1 – S3) but included surprisal of the pre-target content word instead of target surprisal. We compared the relation between neural responses to the pre-target content word and its surprisal under low versus high ambiguity following the same procedure as in the target word analyses (see *Neural responses to surprisal as a function of ambiguity* above and Table S6 for full model structure).

## Results

We constructed German sentences containing one of two polysemic target words: ‘Blatt’ (meaning ‘paper’ or ‘leaf’) or ‘Tor’ (meaning ‘gate’ or ‘goal’; 150 sentences per target). Target words never repeated on consecutive trials to mitigate direct meaning priming. The target word was always placed in the middle of a sentence, preceded by an average of 6 words (range 5 to 9 words). Both target words had balanced meaning frequency (Moritz et al., 2001), which left sentence context the primary source of disambiguation. A native speaker of German trained in linguistics manipulated the pre-target context to bias target interpretation towards one of its meanings to varying extent.

### Surface-based prediction error fails to account for reading times under ambiguity

We first asked whether GPT-2 surprisal, extracted from the model pre-trained on German text corpora, explained reading times for low-and high-ambiguity polysemic words. We used GPT-2 surprisal as a surface-based proxy of prediction error, reflecting how statistically unexpected the polysemic word form is given prior context. Crucially, exposure to statistical regularities in large text corpora allows LLMs, including the early-generation GPT-2 model, to approximate context-specific meaning of polysemic word forms through contextualized embeddings (Coenen et al., 2019; Tripodi, 2021). Therefore, GPT-2 offers a rigorous test of whether a surface-based approximation of meaning suffices to explain human next-word prediction. We hypothesized that the relation between higher GPT-2 surprisal and slower polysemic word reading would persist under ambiguity if prediction is surface-based (Fig. 1c, top), but that it would be attenuated under ambiguity if prediction is meaning-based (Fig. 1c, bottom).

We tested this hypothesis in the online Experiment 1 (Fig. 2a), where participants (*N* = 47–52 per sentence) read each sentence word-by-word in a self-paced manner up to and including the target. After each sentence, participants moved a slider to rate which target meaning was more likely given the preceding context. The average ratings for each sentence validated our context manipulation: they covered both target interpretations and varied continuously from low to high ambiguity (Fig. 2b). For all further analyses, we quantified target ambiguity as the negative absolute rating of target meaning in each sentence.

To test the effects of ambiguity and surprisal on reading times, we fitted a linear mixed-effects regression to log-transformed target reading times (median 301 ms, IQR 152 ms, range 60–2255 ms; Table S1). Reading was slower with higher target ambiguity (*β* ± standard error = 0.03±0.01, *t*(298.4) = 2.31, *p* = 0.02). Likewise, reading was slower with higher surprisal of the preceding word (henceforth, surprisal at lag 1; *β* = 0.03±0.009, *t*(251.1) = 3.15, *p* = 0.002), in line with established spill-over effects (Wilcox et al., 2023). While we found no main effect of target surprisal on reading times (*p* = 0.87), its effect depended on target ambiguity (ambiguity-surprisal interaction *β* = –0.04±0.01, *t*(264.2) = –3.68, *p* < 0.001). To dissect this interaction, we compared the effect of surprisal between sentences with low-and high-ambiguity target words matched on surprisal (AS-orthogonal sentences, 50 sentences per target word; Fig. 2c, Table 1; target reading times under low ambiguity: mean 326 ms, SD 148 ms; under high ambiguity: mean 337 ms, SD 163 ms). In these sentences, the ambiguity-surprisal interaction remained significant (*β* = –0.16±0.05, *t*(88.6) = –3.44, *p* < 0.001; Fig. 2d, Table S2). As expected, under low ambiguity, higher target surprisal led to slower reading (*β* = 0.09±0.03, *t*(88.0) = 2.87, *p* = 0.005). Critically, however, surprisal did not predict reading times under high ambiguity (*β* = –0.07±0.03, *p* = 0.053; note that the effect is marginally significant yet numerically reversed). That is, ambiguity suppressed the ability of a surface-based proxy of prediction error to account for reading times. This supports the meaning-over the surface-based prediction model (Fig. 1c).

**Table 1.** Examples of low-and high-ambiguity target words matched on surprisal.

| Sentence with the target word in bold | Translation | Target GPT-2 surprisal, percentile | Target ambiguity subset |
| --- | --- | --- | --- |
| Die Burg öffnete schon das **Tor**... | The castle had already opened its **gate**... | 5 | low |
| Schwer ist, wieder mehr als zwanzig **Tore**... | It's tough, more than twenty **[goals/gates]** again... | 9 | high |
| Es fielen nun leider nur **Tore**... | Unfortunately, the only **goals** scored... | 87 | low |
| Es sah danach aus als wären hier **Tore**... | It looked like here there were **[gates/goals]**... | 97 | high |

Experiment 1 left several questions open. First, it required participants to make explicit judgments about target meaning. If the task did not incentivize processing of target ambiguity during more naturalistic reading, would the ambiguity-surprisal interaction remain? Second, rich discourse-level context in natural speech rarely leaves word meaning fully ambiguous, which makes sentences with high target ambiguity potentially less plausible. If so, could the effect of target ambiguity be mediated by overall sentence plausibility? We addressed both questions in the online Experiment 2 (Fig. 3a). Participants (= 55) read sentences in a self-paced manner and rated their plausibility by moving a slider. Specifically, participants were instructed to indicate how likely they were to encounter each sentence in an online blog, a book, or in others’ speech. The stimuli were 400 complete grammatical sentences: 100 AS-orthogonal sentences from Experiment 1, 100 semantically plausible filler sentences, and 200 sentences that contained a semantically anomalous word.

**Figure 3.**
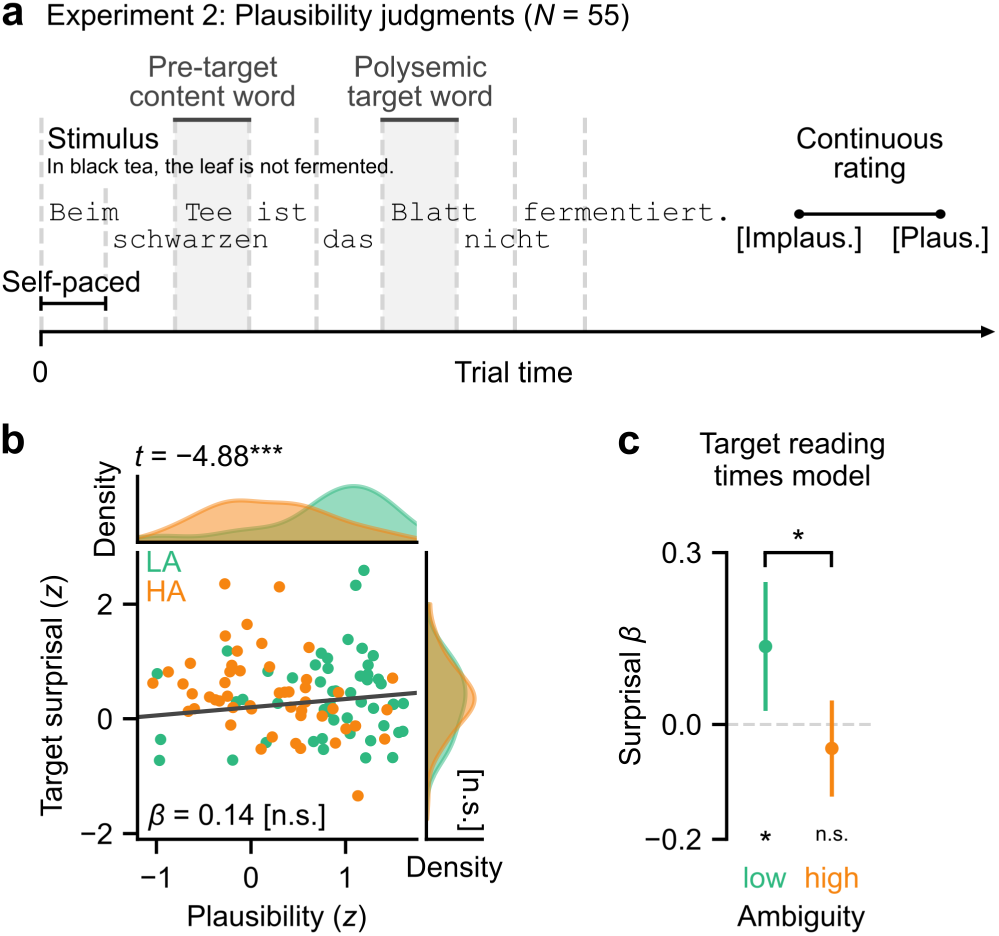
The effect of sentence plausibility on the ambiguity-surprisal interaction. (a) Single trial setup in Experiment 2. Participants read full sentences on the screen word by word in a self-paced manner, a subset of which contained polysemic target words. After the sentence, participants rated its perceived plausibility on a continuous analog scale. (b) The relation between sentence plausibility, target surprisal, and ambiguity (dark grey line shows linear regression fit; adjusted *R^2^* = 0.01). Top and right insets show the distributions of plausibility and surprisal values, respectively, in AS-orthogonal sentences. Plausibility was higher for the low-ambiguity (LA) subset of AS-orthogonal sentences. HA – high ambiguity. (c) *β* coefficients for target surprisal based on a linear mixed-effects regression of log-transformed target reading times. The model contained the same predictors as the model used in Experiment 1, as well as sentence plausibility (based on behavioral ratings, see (a)) and its interactions with target surprisal and ambiguity. GPT-2 surprisal predicted target reading times under low ambiguity but not under high ambiguity, replicating Experiment 1. Error bars represent 95% confidence intervals. * *p* < 0.05, *** *p* < 0.001, n.s. – not significant.

As expected, all AS-orthogonal sentences were rated as more plausible than those with semantically anomalous words (high ambiguity vs. anomalous: *t*(58.3) = 11.1, *p* < 0.001; low ambiguity vs. anomalous: *t*(57.4) = 17.1, *p* < 0.001; Fig. S1). However, sentences with high-ambiguity target words were also rated as less plausible than those with low-ambiguity target words (*t*(97.8) = –4.88, *p* < 0.001; Fig. 3b). To test if the ambiguity-surprisal interaction was mediated by sentence plausibility, we included plausibility and its interactions with target ambiguity and surprisal in the model of target reading times for AS-orthogonal sentences in Experiment 2 (target reading times under low ambiguity: mean 361 ms, SD 167 ms, under high ambiguity: mean 342 ms, SD 146 ms). This analysis replicated the ambiguity-surprisal interaction (*β* = 0.18±0.07, *t*(85.3) = 2.62, *p* = 0.01; Fig. 3c, Table S3). Under low ambiguity, higher target surprisal led to slower reading (*β* = 0.14±0.06, *t*(85.5) = 2.45, *p* = 0.02). However, surprisal did not predict reading times under high ambiguity (*β* = –0.04, *p* = 0.31). Critically, neither the main effect of sentence plausibility nor its interactions with target ambiguity or surprisal were significant (all *p* > 0.28). Overall, Experiment 2 showed that ambiguity suppresses the ability of GPT-2 surprisal to predict reading times independently from sentence plausibility, extending the findings of Experiment 1 to a more naturalistic reading task.

In summary, although GPT-2 surprisal, a surface-based proxy of prediction error, predicted longer reading times for unambiguous words, it failed to predict reading times for high-ambiguity polysemic words matched in surprisal. This pattern was independent of sentence plausibility and extended to a more naturalistic reading task, converging towards the meaning-as opposed to the surface-based prediction model (Fig. 1c).

### A shared cortical network supports ambiguity resolution and prediction error computation

To reveal the cortical dynamics of how ambiguity processing shapes next-word prediction, we recorded MEG while participants (*N* = 27) listened to the sentences of Experiment 1. To ensure attentive listening, participants were occasionally asked to choose a picture that best represented the meaning of the target word in the last sentence. Participants attended to the task: on average, 91.2% (range 75–100%) of their responses matched the sign of the meaning rating obtained in Experiment 1 (Fig. 2b). Because stimulus acoustics correlated with target ambiguity and surprisal (Fig. S2), we regressed acoustically driven responses from neural activity using a spectro-temporal response function (see *Materials and Methods* and Fig. S3) and performed all further analyses on the residual responses, *z*-scored to the interval from –250 to –50 ms before sentence onset.

To isolate the signatures of ambiguity processing from those of prediction error, we compared neural responses to low-and high-ambiguity target words in AS-orthogonal sentences. A spatiotemporal cluster permutation *t*-test revealed two left-and two right-hemispheric clusters; their response profiles were qualitatively similar between the hemispheres (Fig. S4). In the posterior clusters, neural responses were sensitive to ambiguity shortly before target word onset (left-hemispheric cluster in Fig. 4a, cluster < 0.01). Note that the polysemic target words alternated across consecutive trials in our task design, making trial structure strongly predictive of the target word. The pre-onset effect thus suggests that ambiguity of an expected target word is processed prior to its onset in strongly predictive contexts. In the anterior clusters, neural responses were sensitive to ambiguity starting roughly 400 ms post-onset (left-hemispheric cluster in Fig. 4b, cluster *p* < 0.001). Thus, even in strongly predictive contexts where ambiguity *can* be processed even before word onset, it is nonetheless also processed throughout the expected N400 window of prediction error computation (Frank et al., 2015; Heilbron et al., 2022). These temporal dynamics are consistent with a possible interaction between ambiguity processing and prediction error computation, as predicted by the meaning-based model.

**Figure 4.**
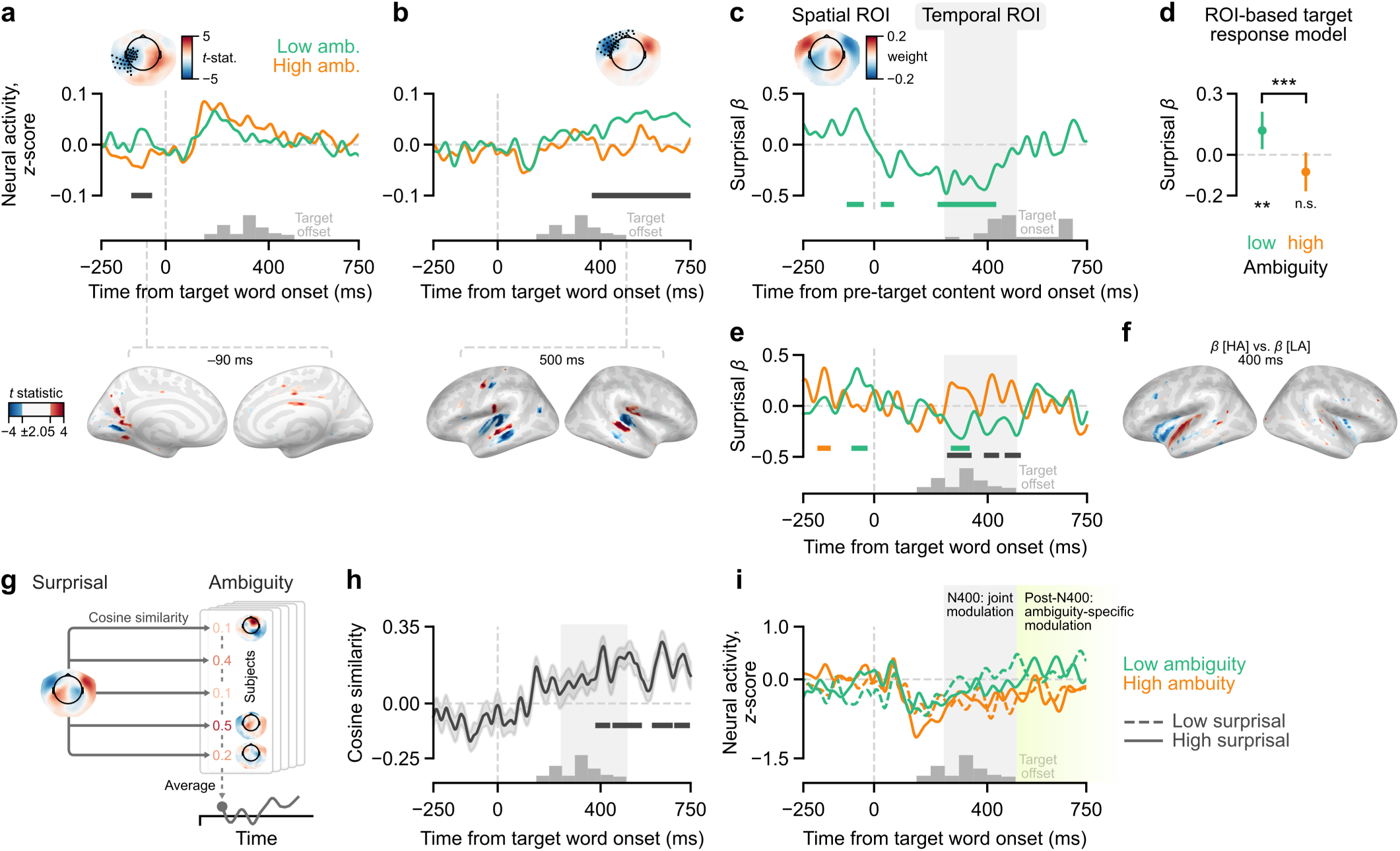
The relation between GPT-2 surprisal, ambiguity, and neural responses. (a) *Top:* Left posterior spatiotemporal cluster of significant response differences for the low-versus high-ambiguity target words, before their onset. The horizontal black line indicates the temporal extent of the cluster. The topography shows the mean *t*-statistic over the cluster; black dots mark the spatial extent of the cluster. *Bottom:* Peak response differences in source space. The *t*-statistic is thresholded at ±2.05 (two-tailed, *α* = 0.05, uncorrected) and is shown for a representative time point. Positive and negative values indicate current direction relative to the cortical surface. (b) Same as (a) for the left anterior spatiotemporal cluster. The topography displays the mean *t*-statistic over a representative interval around 500 ms after target word onset. (c) Region of interest (ROI) definition. The *β* coefficients quantifying the effect of surprisal on neural responses to the pre-target content word under low target ambiguity were subjected to a principal component analysis (PCA; see *Materials and Methods*). The plot shows the weights and the time course of the first PC. The weights were sign-flipped to align all *β* coefficients to the polarity of the left-hemispheric responses in (b). Black horizontal lines indicate significance against zero (cluster-level *α* = 0.05). Grey shading shows the temporal ROI defined *a priori* from 250 to 500 ms post-onset. (d) Target surprisal effects from a linear mixed-effects model of the target word responses, aggregated over the spatiotemporal ROI shown in (c). Surprisal predicted neural responses under low, but not high, ambiguity. Positive values denote stronger effect on neural response magnitude. Error bars represent 95% confidence intervals. ** *p* < 0.01, *** *p* < 0.001. (e) *β* coefficients for target surprisal from a time-resolved linear regression of neural responses to the target word. Coefficients were aggregated over the spatial ROI shown in (c). Horizontal lines indicate significance against zero (in color) or between ambiguity conditions (in black), with cluster-level *α* = 0.05. GPT-2 surprisal had no effect on neural responses to words in the high-ambiguity condition throughout word processing. (f) Peak ambiguity-surprisal interaction in the source space. The *t*-statistic quantifies the difference between surprisal *β* coefficients under low versus high ambiguity. The *t*-statistic is thresholded at ±2.05 (two-tailed, *α* = 0.05, uncorrected) and is shown for a representative time point. Positive and negative values indicate current direction relative to the cortical surface. (g) Cosine similarity was computed between the grand average topography of the surprisal effect (left; the first principal component of surprisal *β* coefficients in the pre-target content word window of low-ambiguity sentences) and participant-specific topographies of the ambiguity effect in the target word window over time (right). (h) Spatial alignment between the effects of surprisal and ambiguity from roughly 390 to 740 ms after target word onset. Horizontal lines indicate significance against zero, cluster-level α = 0.05. (i) Average neural responses to the low-versus high-ambiguity target words, median-split by surprisal. Responses were aggregated over the spatial ROI for surprisal effects shown in (c), motivated by its spatial alignment to the ambiguity effect, as shown in (h). The lime shading illustrates the sustained modulation of neural responses by ambiguity. In (e), (h), (i), the grey shading reproduces the temporal ROI from 250 to 500 ms for illustration purposes.

Next, we aimed to evaluate whether ambiguity suppresses the ability of GPT-2 surprisal to account for neural responses, as expected under the meaning-based prediction model. To maximize the power of this analysis, we focused it on a region-of-interest (ROI). Temporally, the ROI was constrained to 250 to 500 ms after word onset, known as the N400 window where surprisal modulates neural responses (Frank et al., 2015; Heilbron et al., 2022). Spatially, the ROI was defined to capture the largest proportion of variance in the effects of surprisal on neural responses. The definition was based on the pre-target content word window of low-ambiguity sentences (Fig. 2a), which ensured that the ROI was orthogonal to putative effects of surprisal in the target word window. The first principal component of the *β* coefficients for surprisal explained 41% of their variance over time and, crucially, captured the MEG counterpart of the N400 response (Fig. 4c; temporal cluster permutation test, cluster from 230 to 420 ms, *p* < 0.005). We thus used the sensor weights of this first principal component as our spatial ROI.

Within this spatiotemporal ROI, a linear mixed-effects regression of neural responses to the target word revealed a significant ambiguity-surprisal interaction (*β* = –0.21±0.06, *t*(94.0) = –3.40, *p* = 0.001; Fig. 4d, Table S4). Under low ambiguity, higher target surprisal led to stronger neural responses (*β* = – 0.12±0.04, *t*(83.2) = –2.75, *p* = 0.007). However, high ambiguity eliminated the effect of surprisal (*β* = 0.08±0.05, *p* = 0.07, note the numerically opposite direction of the effect). This pattern was consistent between AS-orthogonal and all sentences (Tables S4, S5). It matches our behavioral findings and further supports the meaning-based prediction model.

However, could GPT-2 surprisal account for neural responses even under high ambiguity outside of the N400 window? This would support a combined model where both word meaning and word form co-occurrence statistics contribute to prediction error, as opposed to a strictly meaning-based prediction model. To test this, we analyzed the effects of surprisal on neural responses in a time-resolved fashion. As in the ROI analysis, the magnitude of neural responses increased with target surprisal around 250– 500 ms post-onset for low-ambiguity target words, and this effect was significantly stronger than for high-ambiguity target words (Fig. 4e). Furthermore, we did not find any evidence that GPT-2 surprisal modulated neural responses to high-ambiguity target words at any time point after their onset. Thus, a surface-based proxy of prediction error failed to account for neural responses consistently throughout the processing of high-ambiguity words.

Finally, we investigated the spatio-temporal relation between the effects of target ambiguity and GPT-2 surprisal. We observed a striking spatial alignment between the post-onset effect of target ambiguity and the effect of surprisal: both the sensor-level ROI capturing this effect in the pre-target content word window (topographies in Fig. 4b vs. 4c) and the source-level peak of the ambiguity-surprisal interaction (Fig. 4b vs. 4f). Namely, both effects peaked in the lateral temporal cortex and overlapped in the superior temporal gyrus (Fig. 4b, f). Time-resolved cosine similarity (Fig. 4g) showed that the effects of ambiguity in individual participants spatially aligned with the network generating surprisal responses roughly from 390 to 740 ms post-onset (Fig. 4h; four significant temporal clusters, all *p* < 0.006). In this shared network, the magnitude of neural response concurrently increased with ambiguity or surprisal in the N400 window, but the ambiguity-driven response, in particular, was sustained beyond (Fig. 4i).

In summary, the neural signatures of ambiguity and prediction error processing emerged from a shared cortical network, where ambiguity also suppressed the ability of GPT-2 surprisal to capture the N400 signatures of prediction error. This suggests that ambiguity resolution and prediction error computation are not only mechanistically linked but also spatio-temporally aligned cortical processes.

### Dispersion of surface-based predictions does not account for neural effects of ambiguity

Neural responses are less sensitive to word surprisal if the predictions of that word are uncertain, such that multiple words are expected to a similar degree (Guilleminot and Morillon, 2025). In our study, neutral contexts that render polysemic words ambiguous could as well render GPT-2 predictions of these words uncertain. If so, neural responses might become insensitive to GPT-2 surprisal not because it is an inadequate model of prediction error under ambiguity, but merely because it is derived from highly uncertain model predictions.

To address this, we first examined whether prediction uncertainty increased with ambiguity. We quantified prediction uncertainty at the target word position using the probability distribution over all tokens predicted by GPT-2 at this position. From this distribution, we computed Rényi entropies at the order of 0.1 (henceforth, dispersion, or how many words have comparable above-zero probabilities), 1 (average expected surprisal, or Shannon entropy), and 10 (prediction strength, or how prominent the dominant prediction’s probability is over others; Guilleminot & Morillon, 2025). We also quantified how much the target word shifted the predicted word probability distribution as Kullback-Leibler divergence between GPT-2 predictions after versus before this word. Dispersion increased with ambiguity and was the only metric of prediction uncertainty that explained unique variance in target ambiguity (Fig. 5a). Interestingly, their relation appeared non-linear (Fig. 5b). To formally test this, we fitted a linear regression of ambiguity against dispersion, where both were quantile-transformed, which removed the linearity assumption. This significantly improved model fit (*t*(49) = –10.03, *p* < 0.001; Fig. 5b). The non-linearity reflects that low ambiguity maps to the full range of dispersion values, whereas high ambiguity consistently maps to high dispersion.

**Figure 5.**
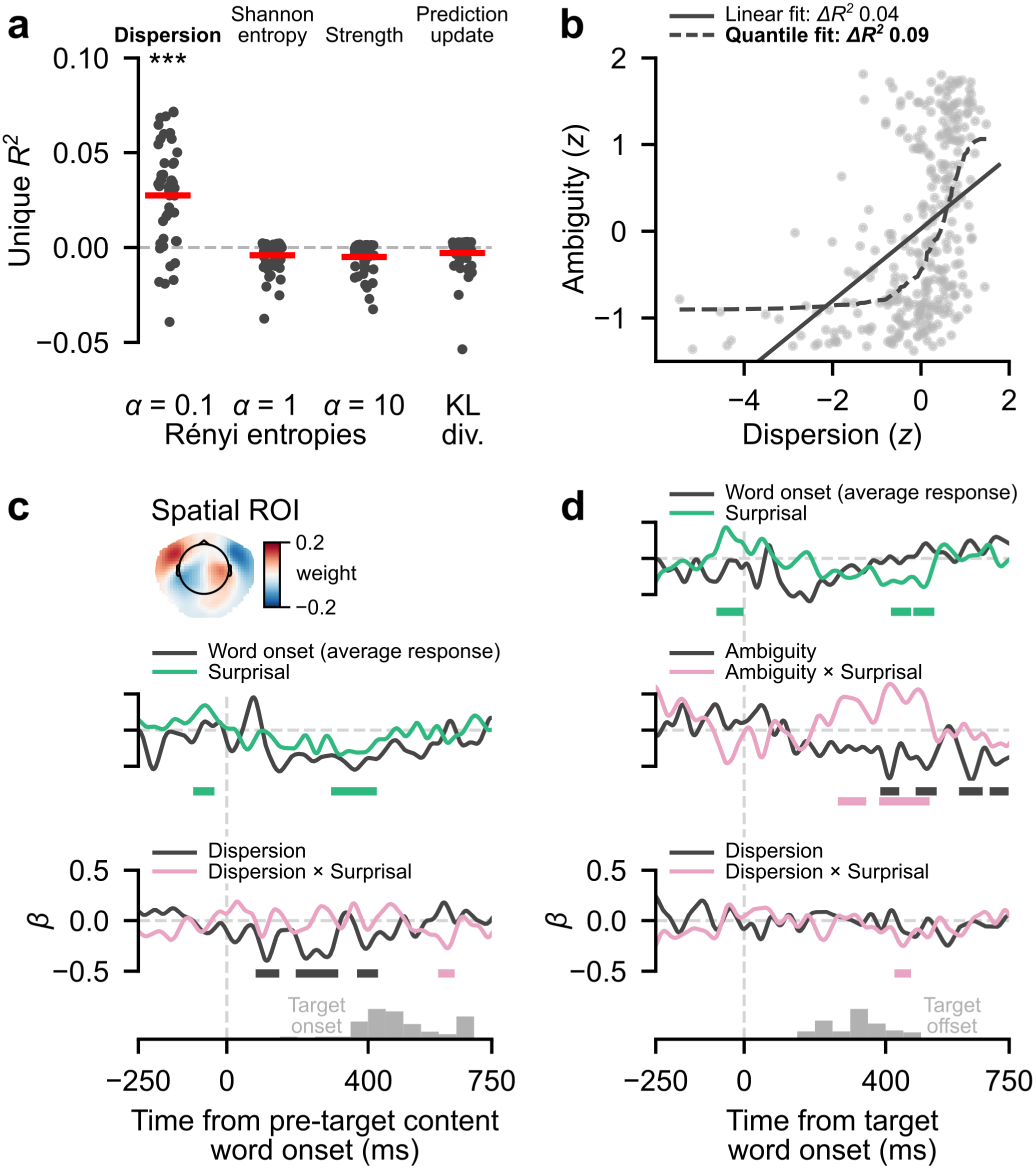
The relation between ambiguity, GPT-2 prediction uncertainty, and neural responses. (a) Unique variance in target ambiguity explained by each uncertainty metric. Only dispersion (Rényi entropy at *α* = 0.1) was related to ambiguity. Asterisks mark the significance of upper-tailed one-sample *t*-tests of these *R^2^* values against zero. KL div. – Kullback-Leibler divergence. * *p* < 0.05, ** *p* < 0.01, *** *p* < 0.001. (b) The non-linear relation between dispersion of GPT-2 predictions at the target word position and target ambiguity across sentences (*N* = 300). Low ambiguity maps to the full range of dispersion values, whereas high ambiguity consistently maps to high dispersion. The quantile fit was obtained by fitting a linear regression to quantile-transformed ambiguity and dispersion values, then projecting its predictions back into the original space. *ΔR^2^* is the unique variance that dispersion explains in target ambiguity above and beyond GPT-2 surprisal. (c) *β* coefficients from a time-resolved linear regression of neural responses to the pre-target content word aggregated over the spatial ROI (topography reproduced from Fig. 4c). The horizontal lines indicate when the *β* coefficients differed significantly from zero (cluster-level *α* = 0.05). Neural responses to the pre-target content word were sensitive to the dispersion of GPT-2 predictions at this word’s position. (d) Same as (c) for neural responses to the target word. Neural responses to the target word were sensitive to its ambiguity, but not to the dispersion of GPT-2 predictions at the target word position. The ambiguity-surprisal interaction was significant with dispersion accounted for.

To test whether dispersion mediated the effects of ambiguity, we fitted a time-resolved linear regression of neural responses to AS-orthogonal sentences against ambiguity, dispersion, and their interactions with surprisal. We conducted this analysis in the same spatial ROI, where we previously found that ambiguity suppressed the effects of target surprisal (Fig. 4c, e). Neural responses to the pre-target content word were sensitive to its dispersion (Fig. 5c; three clusters from 90 to 420 ms, all *p* < 0.01; for full model time courses, see Fig. S5). This replicated recent findings (Guilleminot and Morillon, 2025) and confirmed that the statistical power was sufficient to detect the effects of dispersion in the target word window. However, we did not find any evidence that the target word responses were modulated by dispersion. Furthermore, the interaction between target dispersion and surprisal was brief and only marginally significant (Fig. 5d; one cluster from 440 to 460 ms; *p* = 0.04). Target word responses remained sensitive to ambiguity, as previously (Fig. 5d, main effect of ambiguity: four clusters from 400 to 740 ms, all *p* < 0.05; qualitatively similar to Fig. 4b). Critically, the ambiguity-surprisal interaction also remained significant (Fig. 5d, interaction effect: two clusters from 280 to 510 ms, both *p* < 0.003; qualitatively similar to Fig. 4e). Thus, neural responses became insensitive to GPT-2 surprisal specifically due to ambiguity, not the higher dispersion of GPT-2 predictions in neutral contexts.

In summary, although ambiguity was reflected in the dispersion of surface-based predictions, surprisal of surface-based predictions remained an inadequate model of prediction error under ambiguity even when dispersion was accounted for.

### Prediction error computation is shaped by meaning co-activation, not contextual correlates of ambiguity

Ambiguity is the consequence of polysemy and neutral context combined: a word form with multiple meanings needs to appear in a context sufficiently neutral to induce meaning co-activation. Hence, in a control analysis, we investigated whether a neutral context alone, in the absence of meaning co-activation, would suppress the effect of GPT-2 surprisal. For all three experiments, we tested whether the ambiguity-surprisal interaction would emerge already at the content word immediately preceding the polysemic target word (henceforth, pre-target content word; Fig. 2a, 3a), for which meaning co-activation is minimal. In the behavioral experiments (Tables S1 – S3), higher surprisal at lag 1 tended to slow down reading, although the effect was less robust than for the target word (Exp. 1, full sentence set: *β* = 0.03±0.01, *t*(215.0) = 3.02, *p* = 0.003; Exp. 1, AS-orthogonal sentences: *β* = 0.03±0.016, *t*(74.5) = 1.56, *p* = 0.12; Exp. 2, AS-orthogonal sentences: *β* = 0.04±0.02, *t*(68.8) = 1.81, *p* = 0.07). However, the main effects of surprisal and ambiguity were not significant (all *p* > 0.09), and, critically, neither was their interaction (Exp. 1, full sentence set: *β* = 0.02, *p* = 0.06; Exp. 1, AS-orthogonal sentences: *β* = 0.0002, *p* = 0.99; Exp. 2, AS-orthogonal sentences: *β* = –0.09, *p* = 0.11). That is, the effects of surprisal were generally less robust for the pre-target content word, but were not influenced by context neutrality per se. In the MEG experiment, N400 responses to the pre-target content word did show a main effect of surprisal (*β* = –0.04±0.02, *t*(258.0) = –2.73, *p* = 0.007), but the interaction between surprisal and ambiguity was not significant (*β* = 0.002±0.02, *p* = 0.89). Note that our definition of the spatial ROI maximized surprisal effects at the pre-target content word under low ambiguity, which, if anything, increased the chances of the interaction being significant. In AS-orthogonal sentences (Fig. 7a, Table S6), we found significant main effects of surprisal both under low (*β* = –0.08±0.04, *t*(81.0) = –2.12, *p* = 0.037) and high (*β* = –0.10±0.05, *t*(81.0) = –2.13, *p* = 0.036) ambiguity. In sum, the suppressive effect of ambiguity on the relation between GPT-2 surprisal and prediction error signatures emerged specifically at the polysemic word and was thus driven by the co-activation of multiple word meanings, not context neutrality per se.

## Discussion

The mapping between word surprisal of large language models (LLMs) and neural activity (Frank et al., 2015; Heilbron et al., 2022; Zou et al., 2026) inspires the hypothesis that statistical regularities in word form co-occurrence might underlie predictive language processing in the brain. Here, we provide converging evidence from reading and auditory speech processing against this hypothesis. Specifically, we show that ambiguity disrupts the ability of GPT-2 surprisal to predict reading times and neural responses to speech. This influence of uncertainty in word meaning on prediction error signatures corroborates the essential role of word meaning in predictive language processing.

Our findings extend the emerging suite of linguistic features that attenuate the relation between LLM surprisal and neural responses. Recent work showed that the effect of LLM surprisal on neural responses is weaker for words opening a syntactic chunk than for words continuing a chunk (Zou et al., 2026). Similarly, the effect of LLM surprisal on word reading times is attenuated for ungrammatical compared to grammatical sentences (Slaats et al., 2026). These findings might suggest that human language processing operates upon syntactic features that are not directly encoded by LLMs. However, our findings do not support this syntax-centered view. We describe a lexico-semantic feature, ambiguity, which is independent from syntactic features yet sufficient to entirely abolish the effect of LLM surprisal on behavioral and neural responses. A commonality of syntactically limited or ungrammatical contexts, studied previously, and ambiguous polysemic words, studied here, is that they create uncertainty in sentence meaning. Collectively, this evidence points to an inability of surface-based LLM computations to account for the inference of sentence meaning under uncertainty, regardless of which representational level this uncertainty arises from.

Our findings support the core conceptual assumption of psycholinguistic models that prediction is meaning-based (Rabovsky et al., 2018; Nour Eddine et al., 2024). However, current formulations of these models cannot account for polysemic word prediction, as they are based on a fixed mapping between word forms and semantic features. For instance, polysemic words, such as the word form ‘ball’, would map onto features for both their meanings, including <object>, <round>, <bouncy>, etc., as well as <event>, <festive>, etc. This implies that both sets of meaning features would equally influence prediction error computation, even when prior context clearly favors one meaning. However, our data speak against such an unconstrained prediction error computation. We find that, in strongly disambiguating contexts, polysemic word processing is well explained by LLM surprisal, which is derived from a contextualized representation that approximates the relevant word meaning only (Tripodi, 2021; Wilson & Marantz, 2022). Conceptually, this amounts to a disambiguation process, whose output is then used for prediction error computation. Thus, to account for polysemic word prediction, future models could rely on meaning features instead of word forms, as current psycholinguistic models do, but additionally introduce a disambiguation process to weigh the relevance of word meaning features for computing the prediction error.

While models of predictive language processing and disambiguation have been evolving largely in parallel thus far (Nour Eddine et al., 2022; Rodd, 2020), we provide first evidence that can guide their integration. Our orthogonal manipulation of ambiguity and surprisal within a single MEG experiment concurrently mapped the neural signatures of ambiguity processing and prediction error. Both manifested around the N400 time window and were localized to the lateral temporal cortex. Notably, the effect of ambiguity, but not surprisal, continued at least until 750 milliseconds after the polysemic word onset. A similarly sustained effect of ambiguity has been reported for non-sentential contexts that minimally, if at all, involve prediction (Lee and Federmeier, 2006). We now show that this pattern extends to continuous sentence processing. Overall, these temporal dynamics allow for two possible relations between disambiguation and prediction error computation. Under a *PE-Suspension* hypothesis, disambiguation is an independent computation whose output is a pre-requisite for computing a prediction error. That is, in informative contexts, where disambiguation succeeds, a single prediction error is computed; in neutral contexts, where it fails, prediction error computation is suspended altogether. Alternatively, under a *PE-Maintenance* hypothesis, prediction error is computed for all candidate word meanings, with disambiguation acting as a non-linear function that retains smaller errors and discards larger ones. In neutral contexts, this would result in multiple meaning-specific prediction errors being maintained until further input resolves ambiguity. Notably, a proof-of-principle model implementing a similar mechanism has been recently proposed (Su et al., 2023). Future work is needed to directly contrast it with the *PE-Suspension* architecture and to evaluate it against the temporal dynamics of continuous sentence processing under ambiguity described here.

Lastly, we observed an intriguing effect of polysemy on the fit between the uncertainty, specifically dispersion, in GPT-2 predictions of a word and neural responses to this word. As expected, we found that dispersion modulated neural responses to the pre-target content word, extending recent intracranial findings to non-invasive MEG recordings (Guilleminot and Morillon, 2025). However, dispersion had no effect on neural responses to the polysemic target word. One possibility is that the trial-by-trial predictability of the polysemic target words in our task has led participants to expect these specific words towards the end of the sentence, and thereby made them insensitive to the overall dispersion of the probability distribution. However, another possibility is that GPT-2 dispersion does not account for uncertainty in meaning-specific prediction errors. If so, a *PE-Maintenance* model that differentiates meaning-specific prediction errors should predict neural responses regardless of polysemy. It should also yield dispersion estimates that better predict ambiguity, beyond the moderate relation (9% of shared variance) we found for GPT-2. Follow-up studies are needed to ascertain whether this finding reflects a task-specific prediction strategy or a general effect of polysemy on prediction.

To conclude, we have mapped the spatiotemporal dynamics of disambiguation and prediction error computation in continuous sentence processing. We found that both behavioral and neural signatures of prediction error are shaped by uncertainty in word meaning, a phenomenon that a surface-based prediction mechanism, such as the one implemented in LLMs, cannot capture. Our findings show that prediction error computation relies on word meaning representations and call for the development of computational models that explicate the mechanistic link between disambiguation and predictive language processing.

## Data and Code Availability

The data are currently being analyzed in another project and will be made publicly available upon project completion. The code for analyzing behavioral and pre-processed magnetoencephalography data will be available upon publication at https://github.com/aszyrv/ambiguous_predictions.

## Conflict of Interest Statement

The authors declare no competing financial interests.

## Author Contributions

A.Z.: Conceptualization, Methodology, Investigation, Data Curation, Formal analysis, Visualization, Validation, Writing – Original Draft, Writing – Review & Editing, Project administration, Funding acquisition. V.P.: Investigation, Data Curation. Y.O.: Conceptualization, Methodology, Formal analysis, Visualization, Resources, Writing – Review & Editing, Project administration, Supervision, Funding acquisition.

## Acknowledgements

We thank Gabi Walker-Dietrich, Jürgen Dax, Christoph Braun, and Florian Sandhäger for their help with magnetoencephalography recordings, Joel Baisch for assistance with stimulus development, and all members of the Oganian lab for feedback and discussions throughout the project. The research was supported by Boehringer Ingelheim Fonds (PhD fellowship to A.Z.) and research group funding from the Werner Reichardt Center for Integrative Neuroscience (CIN, to Y.O.).

## Notes

### Competing Interest Statement

The authors have declared no competing interest.

### Summary of Updates

Figure 4 extended with cosine similarity analyses

