## Supplemental Materials for "Semantic ambiguity dissociates the roles of word meaning and co-occurrence statistics in sentence processing"

Andrey Zyryanov[1, 2, 3]*, Victoria Pierz[1], Yulia Oganian[1, 2]*

[1] Centre for Integrative Neuroscience, University of Tübingen, Tübingen, Germany

[2] International Max Planck Research School for the Mechanisms of Mental Function and Dysfunction, Graduate Training Centre of Neuroscience, University of Tübingen, Tübingen, Germany

[3] Max Planck Institute for Biological Cybernetics, Tübingen, Germany

**Table S1. Linear mixed-effects model of log-transformed reading times in Experiment 1 fitted to all sentences.**

| | **Pre-target content word** | | | | | **Polysemic target word** | | | | |
|---|---|---|---|---|---|---|---|---|---|---|
| **Fixed effects** | *β* | *SE* | *t* | *df* | *p* | *β* | *SE* | *t* | *df* | *p* |
| Intercept | -0.11 | 0.06 | -1.67 | 164.10 | 0.097 | -0.07 | 0.08 | -0.94 | 313.34 | 0.350 |
| Sentence position | -0.20 | 0.01 | -13.93 | 209.72 | **<0.001** | -0.21 | 0.02 | -10.12 | 1186.85 | **<0.001** |
| Word position | 0.06 | 0.02 | 2.45 | 271.44 | **0.015** | 0.11 | 0.02 | 4.87 | 277.06 | **<0.001** |
| Length in characters | 0.06 | 0.02 | 3.14 | 282.17 | **0.002** | 0.09 | 0.06 | 1.36 | 279.95 | 0.175 |
| Log-frequency | -0.02 | 0.02 | -1.24 | 223.93 | 0.217 | 0.07 | 0.07 | 1.03 | 285.71 | 0.304 |
| GPT-2 surprisal | -0.01 | 0.01 | -1.10 | 240.00 | 0.274 | 0.00 | 0.01 | 0.17 | 281.05 | 0.868 |
| Target ambiguity | -0.01 | 0.01 | -0.88 | 219.52 | 0.381 | 0.03 | 0.01 | 2.31 | 298.39 | **0.022** |
| GPT-2 surprisal (lag 1) | 0.03 | 0.01 | 3.02 | 214.97 | **0.003** | 0.03 | 0.01 | 3.15 | 251.05 | **0.002** |
| GPT-2 surprisal (lag 2) | 0.00 | 0.01 | 0.10 | 217.12 | 0.924 | -0.00 | 0.01 | -0.18 | 251.25 | 0.860 |
| Sentence × Word position | -0.02 | 0.01 | -1.71 | 12237.15 | 0.087 | -0.01 | 0.01 | -0.47 | 14081.17 | 0.641 |
| Length in characters × Log-frequency | -0.02 | 0.01 | -1.85 | 267.62 | 0.065 | -0.02 | 0.12 | -0.19 | 280.46 | 0.853 |
| GPT-2 surprisal × Target ambiguity | 0.02 | 0.01 | 1.92 | 239.03 | 0.056 | -0.04 | 0.01 | -3.68 | 264.17 | **<0.001** |
| **Random effects** | | | | | | | | | | |
| $\sigma^2$ | 0.34 | | | | | 0.33 | | | | |
| **Random intercept variance** | | | | | | | | | | |
| Item | 0.02 | | | | | 0.01 | | | | |
| Participant | 0.56 | | | | | 0.56 | | | | |
| **Random slopes by participant** | Length in characters × Log-frequency<br>GPT-2 surprisal (lag 2)<br>GPT-2 surprisal (lag 1)<br>Target ambiguity<br>Log-frequency<br>Length in characters<br>Word position<br>Sentence position | | | | | GPT-2 surprisal × Target ambiguity<br>GPT-2 surprisal (lag 2)<br>GPT-2 surprisal (lag 1)<br>Target ambiguity<br>GPT-2 surprisal<br>Log-frequency<br>Word position<br>Sentence position | | | | |
| ICC | 0.0603 | | | | | 0.0506 | | | | |
| N participants | 150 | | | | | 150 | | | | |
| N items | 258 | | | | | 295 | | | | |
| Observations | 12691 | | | | | 14515 | | | | |
| Marginal $R^2$ / Conditional $R^2$ | 0.137 / 0.189 | | | | | 0.130 / 0.174 | | | | |

*Notes.* Target ambiguity was included as a continuous z-scored predictor.

**Table S2. Linear mixed-effects model of log-transformed reading times in Experiment 1 fitted to AS-orthogonal sentences.**

| | Pre-target content word | | | | | Polysemic target word | | | | |
|---|---|---|---|---|---|---|---|---|---|---|
| **Fixed effects** | *β* | *SE* | *t* | *df* | *p* | *β* | *SE* | *t* | *df* | *p* |
| Intercept | -0.15 | 0.07 | -2.24 | 189.13 | **0.026** | 0.05 | 0.11 | 0.41 | 159.74 | 0.680 |
| Sentence position | -0.20 | 0.02 | -10.49 | 309.65 | **<0.001** | -0.25 | 0.03 | -7.49 | 2745.71 | **<0.001** |
| Word position | 0.13 | 0.03 | 4.09 | 76.66 | **<0.001** | 0.11 | 0.04 | 2.72 | 95.65 | **0.008** |
| Length in characters | 0.05 | 0.03 | 2.07 | 99.23 | **0.041** | 0.12 | 0.11 | 1.10 | 87.39 | 0.273 |
| Log-frequency | -0.02 | 0.03 | -0.98 | 74.37 | 0.332 | 0.23 | 0.12 | 1.90 | 87.78 | 0.060 |
| GPT-2 surprisal | -0.01 | 0.03 | -0.42 | 73.52 | 0.677 | -0.07 | 0.03 | -1.96 | 89.87 | 0.053 |
| Target ambiguity [low] | 0.02 | 0.03 | 0.53 | 74.50 | 0.598 | -0.14 | 0.04 | -3.47 | 88.24 | **0.001** |
| GPT-2 surprisal (lag 1) | 0.03 | 0.02 | 1.56 | 74.45 | 0.124 | 0.05 | 0.02 | 3.01 | 88.73 | **0.003** |
| GPT-2 surprisal (lag 2) | 0.01 | 0.02 | 0.59 | 82.61 | 0.560 | -0.00 | 0.02 | -0.10 | 89.05 | 0.919 |
| Sentence × Word position | -0.00 | 0.02 | -0.09 | 3961.56 | 0.928 | 0.02 | 0.02 | 0.89 | 4627.24 | 0.376 |
| Length in characters × Log-frequency | -0.04 | 0.02 | -2.30 | 75.60 | **0.024** | -0.07 | 0.21 | -0.31 | 87.88 | 0.754 |
| GPT-2 surprisal × Target ambiguity [low] | 0.00 | 0.03 | 0.01 | 73.32 | 0.994 | 0.16 | 0.05 | 3.44 | 88.64 | **0.001** |
| **Random effects** | | | | | | | | | | |
| $\sigma^2$ | 0.329 | | | | | 0.338 | | | | |
| **Random intercept variance** | | | | | | | | | | |
| Participant | 0.543 | | | | | 0.573 | | | | |
| Item | 0.010 | | | | | 0.017 | | | | |
| **Random slopes by participant** | Sentence position<br>Word position<br>Length in characters<br>GPT-2 surprisal (lag 2) | | | | | Sentence position<br>Word position<br>GPT-2 surprisal (lag 2) | | | | |
| ICC | 0.627 | | | | | 0.636 | | | | |
| N participants | 150 | | | | | 150 | | | | |
| N items | 87 | | | | | 100 | | | | |
| Observations | 4267 | | | | | 4905 | | | | |
| Marginal $R^2$ / Conditional $R^2$ | 0.061 / 0.650 | | | | | 0.060 / 0.658 | | | | |

*Notes.* The model was fitted to the subset of sentences where the target words differed in ambiguity but matched in surprisal (AS-orthogonal sentences; see Main text, Fig. 2c). Target ambiguity was included as a categorical treatment-coded predictor with high ambiguity as the baseline.

**Table S3. Linear mixed-effects model of log-transformed reading times in Experiment 2.**

| | Pre-target content word | | | | | Polysemic target word | | | | |
|---|---|---|---|---|---|---|---|---|---|---|
| **Fixed effects** | *β* | *SE* | *t* | *df* | *p* | *β* | *SE* | *t* | *df* | *p* |
| Intercept | -0.20 | 0.11 | -1.76 | 69.35 | 0.082 | -0.07 | 0.12 | -0.52 | 101.76 | 0.603 |
| Sentence position | -0.18 | 0.03 | -5.89 | 79.72 | **<0.001** | -0.16 | 0.02 | -6.69 | 49.73 | **<0.001** |
| Word position | 0.07 | 0.07 | 0.91 | 66.11 | 0.364 | -0.05 | 0.08 | -0.62 | 87.04 | 0.540 |
| Length in characters | 0.07 | 0.03 | 1.96 | 91.30 | 0.052 | 0.08 | 0.12 | 0.68 | 85.07 | 0.501 |
| Log-frequency | 0.01 | 0.03 | 0.19 | 69.59 | 0.847 | 0.19 | 0.13 | 1.52 | 88.72 | 0.132 |
| GPT-2 surprisal | 0.01 | 0.03 | 0.40 | 64.31 | 0.690 | -0.04 | 0.04 | -1.02 | 85.15 | 0.309 |
| Target ambiguity [low] | 0.10 | 0.06 | 1.68 | 68.68 | 0.098 | 0.02 | 0.05 | 0.47 | 85.01 | 0.639 |
| Sentence plausibility | -0.02 | 0.05 | -0.31 | 69.55 | 0.756 | -0.02 | 0.05 | -0.42 | 85.03 | 0.679 |
| GPT-2 surprisal (lag 1) | 0.04 | 0.02 | 1.81 | 68.82 | 0.075 | 0.05 | 0.02 | 3.28 | 88.70 | **0.001** |
| GPT-2 surprisal (lag 2) | -0.00 | 0.02 | -0.05 | 69.42 | 0.964 | -0.01 | 0.02 | -0.56 | 85.32 | 0.578 |
| Sentence × Word position | 0.03 | 0.04 | 0.83 | 3881.14 | 0.407 | 0.10 | 0.04 | 2.62 | 56.31 | **0.011** |
| Length in characters × Log-frequency | -0.09 | 0.02 | -3.43 | 70.33 | **0.001** | -0.13 | 0.22 | -0.61 | 85.09 | 0.544 |
| GPT-2 surprisal × Target ambiguity [low] | -0.09 | 0.05 | -1.62 | 67.98 | 0.109 | 0.18 | 0.07 | 2.62 | 85.30 | **0.010** |
| GPT-2 surprisal × Sentence plausibility | 0.06 | 0.05 | 1.26 | 69.95 | 0.212 | -0.00 | 0.07 | -0.05 | 85.20 | 0.959 |
| Target ambiguity [low] × Sentence plausibility | 0.00 | 0.07 | 0.00 | 69.50 | 0.999 | -0.00 | 0.06 | -0.03 | 85.11 | 0.975 |
| GPT-2 surprisal × Target ambiguity [low] × Sentence plausibility | -0.01 | 0.06 | -0.22 | 69.87 | 0.825 | -0.05 | 0.08 | -0.63 | 85.07 | 0.532 |
| **Random effects** | | | | | | | | | | |
| $\sigma^2$ | 0.378 | | | | | 0.315 | | | | |
| **Random intercept variance** | | | | | | | | | | |
| Item | 0.017 | | | | | 0.018 | | | | |
| Participant | 0.524 | | | | | 0.562 | | | | |
| **Random slopes by participant** | GPT-2 surprisal (lag 1)<br>GPT-2 surprisal<br>GPT-2 surprisal × Target ambiguity<br>Target ambiguity<br>Sentence position<br>Word position<br>Length in characters | | | | | Sentence position<br>Word position<br>Sentence position × Word position<br>GPT-2 surprisal (lag 1)<br>Log-frequency | | | | |
| ICC | NA | | | | | 0.055 | | | | |
| N participants | 55 | | | | | 55 | | | | |
| N items | 84 | | | | | 100 | | | | |
| Observations | 4175 | | | | | 4978 | | | | |
| Marginal $R^2$ / Conditional $R^2$ | 0.140 / NA | | | | | 0.100 / 0.150 | | | | |

*Notes.* The model was fitted to the subset of sentences where the target words differed in ambiguity but matched in surprisal (AS-orthogonal sentences; see Main text, Fig. 2c). Target ambiguity was included as a categorical treatment-coded predictor with high ambiguity as the baseline.

**Table S4. Linear mixed-effects model of neural responses to the target word fitted to AS-orthogonal sentences.**

| **Fixed effects** | *β* | *SE* | *t* | *df* | *p* |
|---|---|---|---|---|---|
| Intercept | -0.04 | 0.04 | -1.07 | 57.96 | 0.290 |
| GPT-2 surprisal | 0.08 | 0.05 | 1.85 | 85.82 | 0.067 |
| Target ambiguity [low] | 0.13 | 0.04 | 3.25 | 93.98 | **0.002** |
| GPT-2 surprisal (lag 1) | -0.02 | 0.02 | -0.92 | 93.97 | 0.358 |
| GPT-2 surprisal (lag 2) | -0.03 | 0.02 | -1.36 | 59.77 | 0.179 |
| GPT-2 surprisal × Target ambiguity [low] | -0.20 | 0.06 | -3.40 | 93.98 | **0.001** |
| **Random effects** | | | | | |
| $σ^2$ | 0.910 | | | | |
| **Random intercept variance** | | | | | |
| Item | 0.023 | | | | |
| Participant | 0.024 | | | | |
| **Random slopes by participant, variance** | | | | | |
| GPT-2 surprisal (lag 2) | 0.001 | | | | |
| GPT-2 surprisal | 0.005 | | | | |
| ICC | 0.026 | | | | |
| N participants | 27 | | | | |
| N items | 100 | | | | |
| Observations | 5399 | | | | |
| Marginal $R^2$ / Conditional $R^2$ | 0.010 / 0.036 | | | | |

*Notes.* Single-trial responses for model fitting were averaged over the 250–500 ms interval after the target word onset and over the spatial region of interest (see Main text, Fig. 4c). The model was fitted to the subset of sentences where the target words differed in ambiguity but matched in surprisal (AS-orthogonal sentences; see Main text, Fig. 2c). Target ambiguity was included as a categorical treatment-coded predictor with high ambiguity as the baseline.

**Table S5. Linear mixed-effects model of neural responses to the target word fitted to all sentences.**

| **Fixed effects** | *β* | *SE* | *t* | *df* | *p* |
|---|---|---|---|---|---|
| Intercept | -0.00 | 0.04 | -0.06 | 31.62 | 0.949 |
| GPT-2 surprisal | -0.03 | 0.01 | -2.26 | 134.26 | **0.025** |
| Target ambiguity | -0.04 | 0.01 | -2.85 | 128.94 | **0.005** |
| GPT-2 surprisal (lag 1) | -0.01 | 0.01 | -1.27 | 293.86 | 0.207 |
| GPT-2 surprisal (lag 2) | -0.01 | 0.01 | -0.67 | 97.62 | 0.502 |
| GPT-2 surprisal × Target ambiguity | 0.04 | 0.02 | 2.93 | 75.44 | **0.004** |
| **Random effects** | | | | | |
| $\sigma^2$ | 0.932 | | | | |
| **Random intercept variance** | | | | | |
| Item | 0.023 | | | | |
| Participant | 0.028 | | | | |
| **Random slopes by participant, variance** | | | | | |
| GPT-2 surprisal | 0.000 | | | | |
| Target ambiguity | 0.000 | | | | |
| GPT-2 surprisal × Target ambiguity | 0.002 | | | | |
| GPT-2 surprisal (lag 2) | 0.000 | | | | |
| ICC | 0.026 | | | | |
| N participants | 27 | | | | |
| N items | 300 | | | | |
| Observations | 16194 | | | | |
| Marginal $R^2$ / Conditional $R^2$ | 0.007 / 0.032 | | | | |

*Notes.* Single-trial responses for model fitting were averaged over the 250–500 ms interval after the target word onset and over the spatial region of interest (see Main text, Fig. 4c). The model was fitted to all sentences with target ambiguity as a continuous z-scored predictor.

**Table S6. Linear mixed-effects model of neural responses to the pre-target content word fitted to AS-orthogonal sentences.**

| **Fixed effects** | *β* | *SE* | *t* | *df* | *p* |
|---|---|---|---|---|---|
| Intercept | 0.06 | 0.05 | 1.11 | 74.19 | 0.272 |
| GPT-2 surprisal | -0.10 | 0.05 | -2.13 | 80.96 | **0.036** |
| Target ambiguity [low] | -0.04 | 0.06 | -0.69 | 80.97 | 0.492 |
| GPT-2 surprisal (lag 1) | 0.02 | 0.03 | 0.55 | 80.97 | 0.586 |
| GPT-2 surprisal (lag 2) | -0.01 | 0.03 | -0.43 | 68.02 | 0.669 |
| GPT-2 surprisal × Target ambiguity [low] | 0.01 | 0.06 | 0.23 | 80.97 | 0.818 |
| **Random effects** | | | | | |
| $\sigma^2$ | 0.913 | | | | |
| **Random intercept variance** | | | | | |
| Item | 0.045 | | | | |
| Participant | 0.028 | | | | |
| **Random slopes by participant** | GPT-2 surprisal (lag 2) | | | | |
| ICC | 0.048 | | | | |
| N participants | 27 | | | | |
| N items | 87 | | | | |
| Observations | 4697 | | | | |
| Marginal $R^2$ / Conditional $R^2$ | 0.007 / 0.054 | | | | |

*Notes.* Single-trial responses for model fitting were averaged over the 250–500 ms interval after the pre-target content word onset and over the spatial region of interest (see Main text, Fig. 4c). The model was fitted to the subset of sentences where the target words differed in ambiguity but matched in surprisal (AS-orthogonal sentences; see Main text, Fig. 2c). Target ambiguity was included as a categorical treatment-coded predictor with high ambiguity as the baseline.

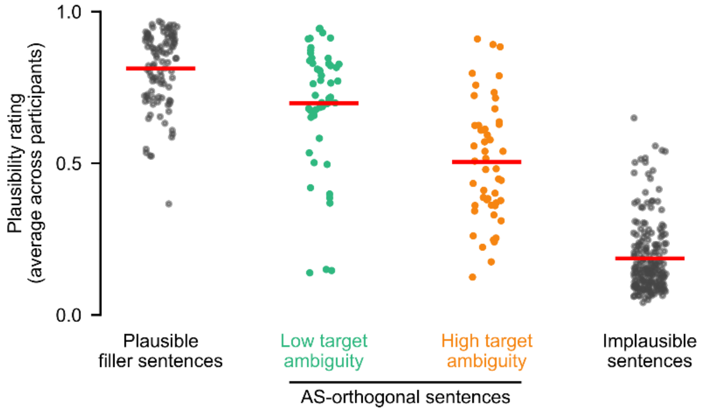


**Figure S1. Plausibility ratings obtained in Experiment 2.** Each observation corresponds to one sentence's rating averaged across 55 participants. The minimum and the maximum of the plausibility scale correspond to the 'sehr unwahrscheinlich' (very unlikely) and 'sehr wahrscheinlich' (very likely) labels shown to the participants. All pairwise differences between conditions are significant (all $p <$ 0.001). AS-orthogonal sentences are those where target ambiguity and surprisal were orthogonalized (see Main text, Fig. 2c).

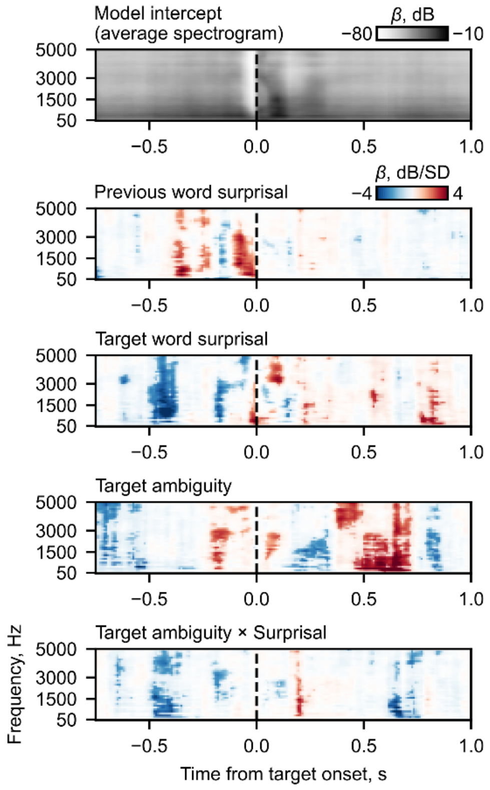


**Figure S2. The relation between target word features and stimulus acoustics in the MEG stimulus set.** The plot shows $\beta$ coefficients from a linear regression of power in each stimulus spectrogram bin against the target word features. The spectrograms ($N$ = 300) were computed using a mel-spaced filter bank with 80 center frequencies spanning 10 – 5000 Hz and segmented from –0.75 to 1 second around the target word onset. $\beta$ coefficients significant at $\alpha$ = 0.05 (uncorrected) are shown in brighter colors.

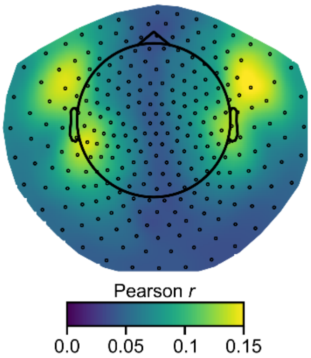


**Figure S3. Topography of the spectro-temporal response function model performance.** The plot shows the training set correlation (average across 27 participants) between the true neural responses and those predicted by a spectro-temporal response function model (see *Materials and Methods*). Note that we did not conduct a full cross-validation procedure because the model was used to regress as much acoustics-related variance in neural responses as possible, rather than to obtain an unbiased estimate of model performance.

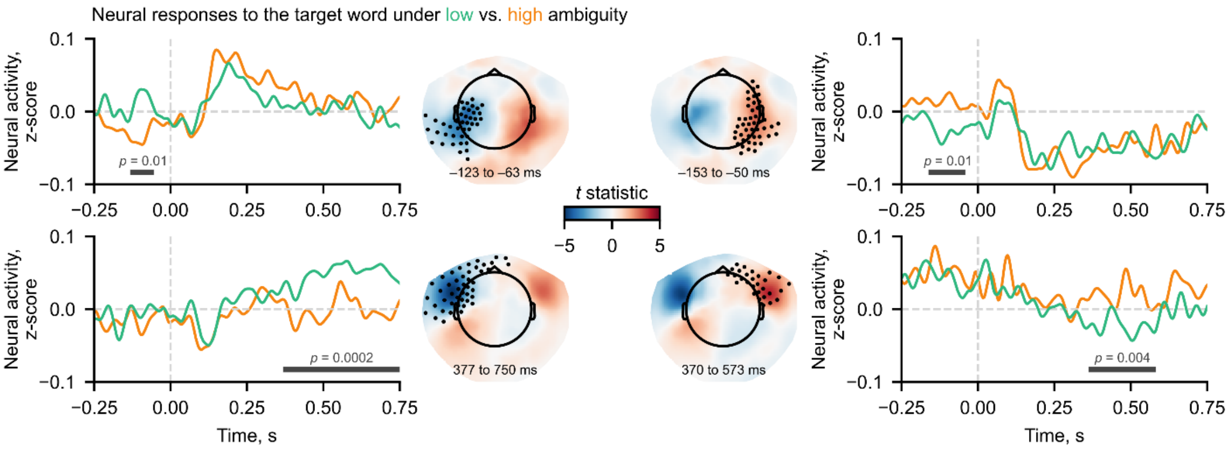


**Figure S4. Spatiotemporal clusters of significant differences in neural responses to high- versus low-ambiguity target words.** The horizontal line indicates the temporal extent of each cluster. The topography displays the mean *t*-statistic over this window. Black dots mark all sensors that belonged to the cluster.

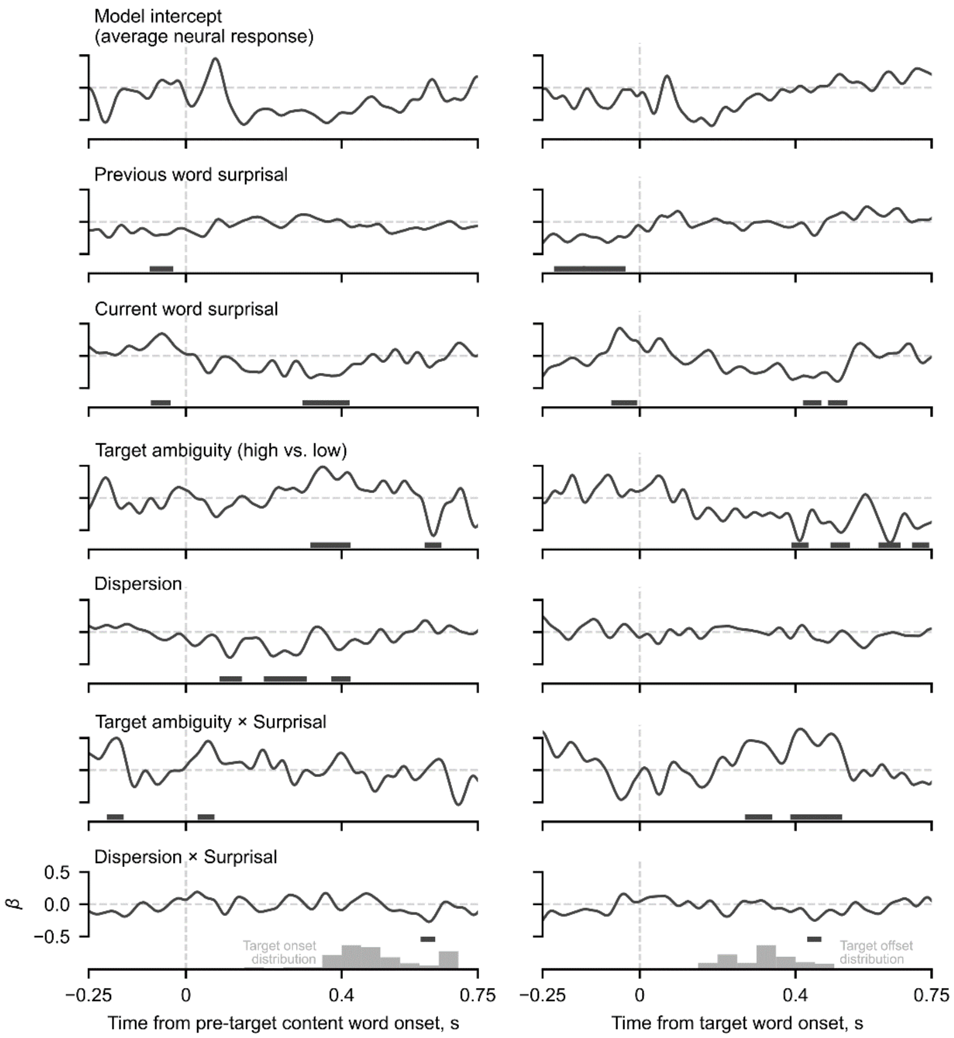


**Figure S5. Linear regression of neural responses against ambiguity, dispersion, and their interactions with surprisal.** Time-resolved $\beta$ coefficients showing the effect of each feature on the neural responses to the pre-target content word (left) and to the target word (right). The coefficients were aggregated over the spatial region of interest shown in Fig. 4c (see Main text). The horizontal lines indicate when the $\beta$ coefficients differed significantly from zero (cluster-level $\alpha$ = 0.05).
